# Live-cell imaging of marked chromosome regions reveals dynamics of mitotic chromosome resolution and compaction

**DOI:** 10.1101/305391

**Authors:** John K. Eykelenboom, Marek Gierliński, Zuojun Yue, Nadia Hegarat, Hilary Pollard, Tatsuo Fukagawa, Helfrid Hochegger, Tomoyuki U. Tanaka

## Abstract

When human cells enter mitosis, chromosomes undergo substantial changes in their organisation to resolve sister chromatids and compact chromosomes. Despite the fundamental importance of this phenomenon to genome stability, we still do not fully comprehend the timing and coordination of these events. To address these questions, we need to evaluate the progression of both sister chromatid resolution and chromosome compaction in one assay. We achieved this by analysing changes in configuration of marked chromosome regions over time, with high spatial and temporal resolution. This assay showed that sister chromatid resolution is an iterative process that begins in late G2 phase and completes in prophase. Cohesins and WAPL antagonistically regulate sister chromatid resolution in late G2 and prophase whilst local enrichment of cohesin on chromosomes prevents precocious sister chromatid resolution. Moreover, our assay allowed quantitative evaluation of the timing and efficiency of condensin II and I activities in promoting sister chromatid resolution and chromosome compaction, respectively. Thus, our real-time assay sheds new light on the dynamics of mitotic chromosome resolution and compaction.

## Introduction

During the cell cycle, chromosomes undergo dynamic changes in their organisation. In interphase, chromosomes remain de-condensed and DNA replication generates sister chromatids (Bickmore and van Steensel, 2013). At the beginning of mitosis, chromosomes undergo two major structural changes in metazoan cells: First, sister chromatids are resolved from each other along chromosome arms; this process involves removal of sister chromatid cohesion and elimination of topological DNA links (Nasmyth and Haering, 2009; Pommier et al., 2016; Uhlmann, 2016). Second, each sister chromatid is compacted; as a result, they become thicker in width and shorter in length (Hirano, 2016; Uhlmann, 2016). These two changes are a prerequisite for proper chromosome segregation towards opposite spindle poles during the subsequent anaphase. However, the precise timing and coordination of these two changes of chromosomes – sister chromatid resolution and chromosome compaction – has so far not been sufficiently analysed.

Several factors regulate sister chromatid resolution and chromosome compaction. Sister chromatids are held together by the cohesin complex, which forms a ring structure consisting of SMC1, SMC3, RAD21 and SA1/2 (Nasmyth and Haering, 2009). For sister chromatid resolution, the cohesin complex must be removed through the destabilising activity of the WAPL along chromosome arms during prophase (prophase pathway) whilst it is retained at the centromere to maintain sister chromatid cohesion until anaphase onset (Morales and Losada, 2018; Peters et al., 2008). In addition, topological DNA links (DNA catenation) that stem from DNA supercoiling during DNA replication, must also be removed by the de-catenation activity of topoisomerase II (topo II) (Piskadlo and Oliveira, 2017; Pommier et al., 2016). It remains unknown at what time in early mitosis cohesins, WAPL and topo II regulate sister chromatid resolution and how they coordinate this process.

Moreover, the condensin complex plays important roles in both sister chromatid resolution and chromosome compaction. The condensin complex exists as two forms – condensin I and II – that consist of the common SMC2 and SMC4 subunits and distinct non-SMC subunits such as NCAPD2 and NCAPD3 (for condensin I and II, respectively) (Hirano, 2012). Evidence suggests that condensin I and II collaboratively generate helical arrays of nested chromatin loops (Gibcus et al., 2018). Furthermore, condensin II operates earlier and contributes more to sister chromatid resolution than does condensin I (Hirano, 2012; Nagasaka et al., 2016). The precise timing of condensin I and II activity and their relative contribution to sister chromatid resolution and chromosome compaction remains to be fully elucidated.

The analysis of chromosome re-organisation in early mitosis has been advanced by several new methods, which include chromosome conformation capture analyses (Hi-C) (Gibcus et al., 2018; Naumova et al., 2013), differential visualisation of sister chromatids (Nagasaka et al., 2016), and in vitro reconstitution of mitotic chromosomes (Shintomi et al., 2015). However, currently available methods still fall short in attaining the following two goals: First, very few methods allow quantitative evaluation of sister chromatid resolution and chromosome compaction together. For example, Hi-C provides detailed information about chromosome compaction, but not about sister chromatid resolution. A simultaneous evaluation of resolution and compaction is, however, critical since these processes are likely to be highly coordinated. Second, although progression of global chromosome re-organisation has been analysed in early mitosis, real-time analysis of regional chromosome re-organisation has so far not been achieved. Since global chromosome changes are the ensemble outcome of regional changes, it could obscure dynamic regional changes of chromosomes – e.g. any rapid or cyclical changes.

To achieve real-time measurements of regional compaction and resolution we decided to investigate changes of specific chromosome regions with time. Using bacteria-derived operator arrays (Belmont and Straight, 1998; Michaelis et al., 1997) we have created a fluorescence reporter system that quantitatively evaluates both sister chromatid resolution and chromosome compaction at chosen chromosome regions in human cells. This has allowed us to study dynamic chromosome reorganisation from G2 phase to early mitosis, by live cell microscopy. By assessing the roles of different factors on these events we reveal how sister chromatid resolution and chromosome compaction are temporally coordinated and how these processes are regulated.

## Results

### Visualising sister chromatid resolution and compaction at a chosen region in live human cells

In order to analyse mitotic chromosome re-organisation, we developed an assay system to visualise sister chromatid resolution as well as chromosome compaction in live HT-1080 diploid human cells. Using CRISPR-Cas9 technology we integrated a *tet* operator array and a *lac* operator array (Lau et al., 2003) with 250 kbp interval, to a region of chromosome 5 with low gene density (Figure 1A; S1A and B). The *tet* and *lac* operators were bound by Tet-repressor fused to four monomer-Cherry fluorescent proteins (TetR-4xmCh) and by the Lac-repressor fused to enhanced green fluorescent protein and a nuclear localisation signal (EGFP-LacI-NLS), thus visualised as red and green fluorescent dots in the cell nucleus (Figure 1B). We chose a cell line where red and green fluorescent dots were found in proximity; reasoning that, in this cell line, *tet* and *lac* operator arrays were integrated on the same copy of chromosome 5 rather than separately on the two homologous chromosome 5s. The *tet* and *lac* operator arrays were stably maintained during cell proliferation since their signal intensity did not become weakened. As implied previously (Chubb et al., 2002; Thomson et al., 2004), integration of these operators onto the chromosome 5 did not affect cell cycle progression or fidelity of chromosome segregation; for example, there was no change in DNA content of these cells as determined by flow cytometry or no missegregation of chromosome 5 observed by microscopy.

**Figure 1:**
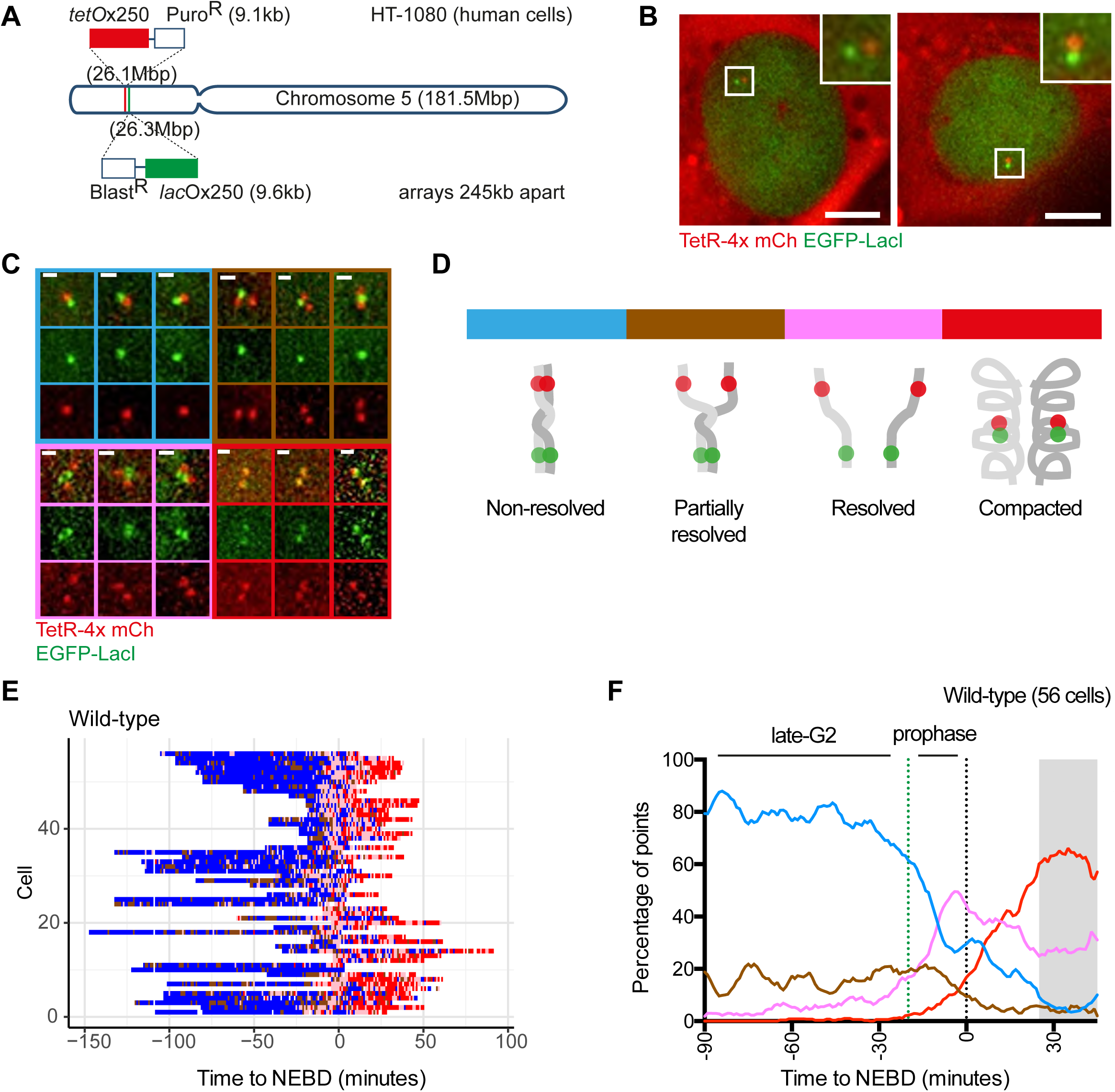
A fluorescent reporter for observing configuration of a selected chromosome region in live cells. (A) Diagram depicting the location of *tet* and *lac* operator arrays introduced into human chromosome 5 using CRISPR-Cas9 gene targeting. Confirmation of targeting is shown in Figure S1(A) and (B). (B) Visualisation of these arrays by expression of TetR-4x mCherry and EGFP-LacI in HT-1080 cells (TT75). White square box represents the zoomed region shown in the upper right-hand corner. Scale bar represents 10µm. (C) Representative images of the major configurations of the fluorescent reporter observed in TT75 cells. Designated colour-codes for each are indicated in the image frames. Scale bar represents 1µm. (D) Diagram representation of the reporter configurations shown in (C). (E) Behaviour of the fluorescent reporter as observed in individual wild-type live-cells plotted across the Y-axis. TT75 cells were arrested by a double thymidine block and subsequently released from the block. 8–10 hr after the release, images were taken every minute and, at each time point, the configuration of the reporter was determined and represented using the colour-codes depicted in (C) and (D). Data from individual cells was aligned according to the occurrence of nuclear envelope breakdown (NEBD; time zero); an example of NEBD is shown in Figure S1(C). White spaces represent time points where configuration could not be determined. (F) The proportion of each configuration was determined from the data shown in (E) with smoothing applied by calculating the rolling mean proportion across 9 minutes. G2 phase and prophase are as indicated. The proportion plots are colour coded according to (C) and (D). In the grey shaded area, the proportion was obtained from less than 10 cell samples at each time point, and is therefore less reliable. See also Figure S1.

Imaging the above cells revealed various configurations of the fluorescent dots. During interphase, which makes up the majority of the cell cycle, we observed one green dot and one red dot (Figure 1C, defined as blue state). However, in early mitosis (shortly before and after nuclear envelope breakdown [NEBD]; see below), we often observed i) one green dot and two red dots (or vice versa; defined as brown state), ii) two green dots and two red dots without co-localisation (defined as pink state) and iii) two green dots and two red dots with colocalisation of each green and red dot (defined as red state). These states are likely to reflect chromosome re-organisation during early mitosis, as follows (Figure 1D): a) the blue state represents ‘non-resolved’ sister chromatids (if cells have progressed through S phase), b) the brown state reflects ‘partially resolved’ sister chromatids, c) the pink state shows ‘resolved’ (but not compacted) sister chromatids, and d) the red state indicates resolved and ‘compacted’ chromatids.

To analyse chromosome re-organisation in early mitosis, we released cells with the fluorescent dots from a double thymidine block, and acquired live-cell images every minute between 8 hr and 12 hr (relative to the release). We were able to identify the timing of NEBD in individual cells, as it caused dispersion of the EGFP-LacI-NLS signal (specifically the fraction not bound to *lac* operators) from the nucleus to the whole cell (Figure S1C). For individual cells, we aligned the sequence of the states of fluorescent dots (as defined in Figure 1C and D), relative to NEBD (defined as time zero) (Figure 1E). Then we plotted the proportion of cells displaying each state, against time (Figure 1F). We noticed that as cells approached NEBD, the pink ‘resolved’ state increased its frequency. After NEBD, there was an increase in occurrence of the red ‘compacted‘ state. These observations in unperturbed live cells suggest that chromosome re-organisation proceeds from sister chromatid resolution to chromosome compaction, as assumed in Figure 1D.

To analyse the dynamics of a marked chromosome region in different cells, we inserted *tet* and *lac* operator arrays with 100kbp interval, on the Z chromosome of DT40 cells (Figure S1D and E) and visualised them using the same method as above. In these cells, we could identify the same four configurations of the fluorescent dots as above (Figure S1F). Live-cell imaging revealed appearance of pink ‘resolved’ and red ‘compacted’ states with similar timing to that observed in human cells (Figure S1G). We conclude that a marked chromosome region behaves similarly during early mitosis in different vertebrate species and in different chromosome contexts.

### Sister chromatid resolution is an iterative process that begins in late G2 phase and completes during prophase

Further analysis of the HT-1080 cells revealed that the brown ‘partially resolved’ state often (~20% of time points) appeared up to two hours prior to NEBD (Figure 1E, F). The brown state typically appeared and continued for a few minutes before returning to the blue ‘non-resolved’ state (Figure 2A). Thus, it seems that the blue and brown states show cyclical exchanges before being converted to the pink ‘resolved’ state (Figure 2B).

**Figure 2:**
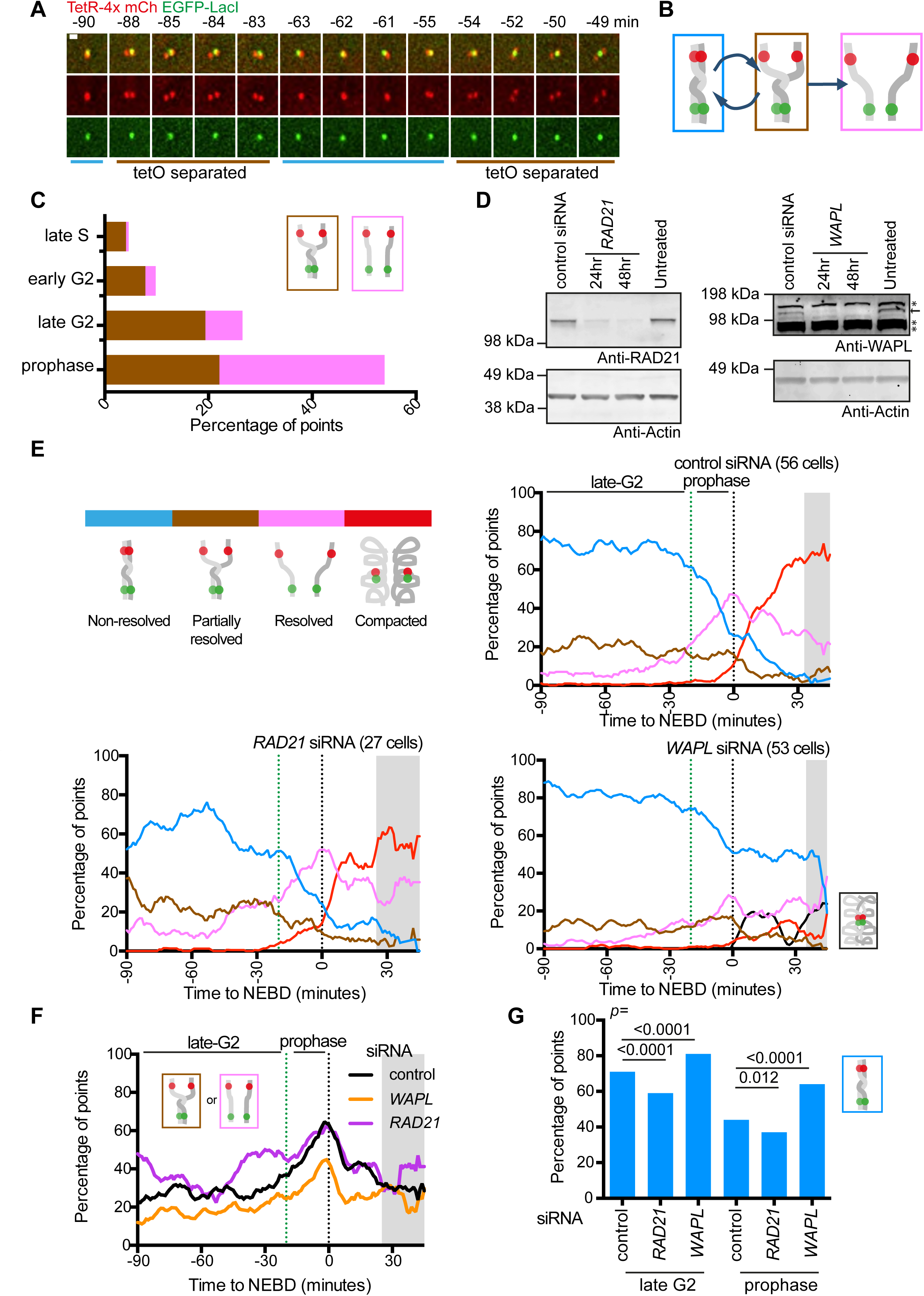
Sister chromatid resolution is an iterative process, which begins in late G2-phase and is antagonistically regulated by cohesins and WAPL. (A) Time lapse images of the fluorescent reporter of an individual cell showing cyclical sister *tet* operator array separation. Times indicated are relative to nuclear envelope breakdown (NEBD). Scale bar represents 1µm. (B) Diagram depicting cyclical sister separation of the *tet* operator array in late G2 phase. (C) The proportion of the brown “partially resolved” or the pink “resolved” state during the indicated cell cycle phases. The cell cycle phases were determined, as explained in text, by visualising PCNA and NEBD in TT104 cells. TT104 cells were generated by integrating Cerulean-PCNA into TT75 cells (see Figure 1(B)). Measurements are taken from 61 (late S and early G2 phase) and 26 (late G2 and prophase) cells. See also Figure S2(A), (B), (C) and (D). The numbers of analysed points were 973, 2713, 768 and 267 for late S phase, early G2, late G2 and prophase respectively. (D) Western blotting analysis of RAD21 (left) or WAPL (right) proteins following treatment with siRNA for the indicated time. he non-specific siRNA treatment was over 48 hours. Probing of the same membranes with an antibody against actin serves as a loading control. The asterisks indicate non-specific binding of the WAPL antibody. (E) The proportion of each configuration in cells treated with control, RAD21 and WAPL siRNA. Reporter configuration is shown by colour codes in diagram (top, left). TT75 cells were treated with siRNA, arrested by a double thymidine block, released from the block and observed by live-cell microscopy, as shown in Figure S2 (E). WAPL siRNA led to co-localisation of all four fluorescent dots (black) after NEBD in some cells, which we interpret as a “non-resolved and compacted” state. Data from individual cells is shown in Figure S2(G), (H) and (I). Smoothing was applied as in Figure 1(F). The start of prophase is indicated by the green dotted line (−20 min) and NEBD by the black dotted line (0 min). The grey shaded area is as Figure 1(F). (F) The proportion of the brown “partially resolved” or pink “resolved” state for control siRNA, RAD21 depleted or WAPL depleted cells. These data were extracted from Figure 2 (E). The end of G2-phase and the start of prophase are indicated as in (E). The grey shaded area indicates a time window where the proportion was obtained from less than 10 cell samples at each time point, at least in one siRNA condition. (G) Graph shows the proportion of the blue “non-resolved” state for control, RAD21 and WAPL siRNA during late G2-phase or prophase. These data were extracted from Figure 2 (E). The indicated two siRNA conditions were compared and *p* values were obtained, by the *chi*-square statistical test, and the number of analysed points were 2457, 1502 and 3081 for control, RAD21and WAPL during late G2 phase, respectively, and 915, 520 and 934 for control, RAD21 and WAPL during prophase, respectively. See also Figure S2.

This raised the possibility that sister chromatid resolution may already begin in G2 phase. To address this, we defined S, G2 and prophase in our real-time imaging, as follows: We identified S-phase cells by visualising a component of the replication machinery, Proliferating Cell Nuclear Antigen (PCNA) tagged with mCerulean. Since Cerulean-PCNA shows characteristic globular signals during S phase (Figure S2A) (Kitamura et al., 2006; Thomson et al., 2010), we defined the end of S phase as the time its globular signals disappeared (Figure S2B, left). Our observations of PCNA and NEBD by live-cell imaging suggested that the length of G2 phase (between the end of S phase and the start of prophase) was 5–7 hrs (Figure S2B, right), which is consistent with conclusions in other studies (Defendi and Manson, 1963). We also defined prophase as a 20-min time window prior to NEBD, according to previous estimates (Liang et al., 2015) and also based on the global morphological change of the chromosomes observed in our cells (Figure S2C).

The brown ‘partially resolved’ state appeared infrequently (4 – 8% of cells) in late S-phase (the last 30 min of S phase) and early G2 phase (first 90 min of G2 phase), but its frequency increased to 19% in late G2 (last 120 min of G2 phase) (Figure 2C, S2D). Subsequently, in prophase, the pink ‘resolved’ state increased to 32% (from 7% in late G2 phase; Figure 2C). The brown state was also observed in up to 20% of DT40 cells in late G2 phase (Figure S1F, G). We conclude that sister chromatid resolution begins in late G2 phase and completes during prophase. Sister chromatid resolution is an iterative process. i.e. a partially-resolved state repeatedly appears in late G2 phase, which may reflect ‘attempts’ at sister chromatid resolution of that region, finally leading to complete sister chromatid resolution in prophase. It has been suggested that sister chromatid resolution begins in early prophase (Nagasaka et al., 2016), but our results imply it begins even earlier. Perhaps, the temporary appearance of partially resolved sister chromatids is discernible only with focused analysis of a small chromosome region, but not by global chromosome analysis.

### Sister chromatid resolution is antagonistically regulated by cohesins and WAPL, not only during prophase but also in G2 phase

Previous studies suggested that maintenance of catenated DNA requires cohesins (Farcas et al., 2011; Sen et al., 2016) and cohesin removal from chromosome arms by WAPL promotes sister chromatid resolution in prophase (Peters et al., 2008). We tested if these conclusions are reproduced in prophase with our assay. We also addressed, if the initial sister chromatid resolution in late G2 phase, described above, depends on cohesin removal by WAPL. We used siRNA to deplete the cohesin subunit RAD21 or WAPL within 24–48 hours of transfection (see Figure S2E and 2D). By observing fixed metaphase chromosomes, we found that sister chromatids were morphologically less distinct after WAPL depletion confirming the previously observed global defect in sister chromatid resolution caused by WAPL siRNA (Gandhi et al., 2006) (Figure S2F).

To assess the roles of cohesin and WAPL on resolution and compaction, we scored how fluorescent dots changed their configuration from late G2 to prometaphase after 48 hours of siRNA treatment (Figure 2E; Figure S2G, H and I). The treatment of cells with control siRNA did not cause any significant changes, compared with untreated cells, in the dynamic behaviour of fluorescent dots (compare Figure 2E, upper right with Figure 1F). However, depletion of RAD21 caused i) an increase of the brown ‘partially-resolved’ and pink ‘resolved’ states (Figure 2F) and conversely, ii) a decrease of the blue ‘non-resolved’ state (Figure 2G), in late G2 and prophase. In contrast, depletion of WAPL led to the opposite outcomes, i.e. i) a decrease of the brown and pink states (Figure 2F) and ii) an increase of the blue state (Figure 2G), in late G2 and prophase. We conclude that cohesins inhibit precocious sister chromatid resolution in late G2 and prophase while WAPL promotes sister chromatid resolution during these phases. Thus, cohesins and WAPL play antagonistic roles in sister chromatid resolution not only during prophase but also during late G2 phase.

### Local reduction of cohesin levels cause precocious sister chromatid resolution during prophase

While we were observing the ‘partially resolved’ configuration of fluorescent dots (the brown state in Figure 1C, D), we found that the *tet* operator array (red fluorescent dot) showed much more frequent sister separation than does the *lac* operator array (green fluorescent dot) (Figure 3A). It is unlikely such sister separation of the *tet* operator array was an artefact of this array, since the *lac* operator array also showed more frequent sister separation (than in the original cell line) when it was integrated at another chromosome region in another cell line (Figure S3A and B). This raised the possibility that, in the original cell line, sister separation of the *lac* operator array was infrequent because of a specific chromosome feature.

We inspected the genomic region where the operator arrays were integrated in the original cell line, using the human genome browser at UCSC (Kent et al., 2002) and publicly available Chromatin Immunoprecipitation sequencing (ChIP seq) data sets therein (ENCODE Project Consortium, 2012). We noticed that the *lac* operator array was located within 5 kbp of a cohesin-enriched region (Figure 3B, SMC3 and RAD21). We thought that this close proximity to a cohesin enriched site might contribute to the reduced (or delayed) sister separation of the *lac* operator array. Further inspection showed that this cohesin peak coincides with a CTCF-enriched region (Figure 3B, CTCF). CTCF is a protein that binds specific DNA sequences and acts as a barrier to cohesin movement leading to its local accumulation (Parelho et al., 2008; Wendt et al., 2008). Therefore, deletion of the CTCF binding sequence of this region might reduce the level of cohesins found there.

**Figure 3:**
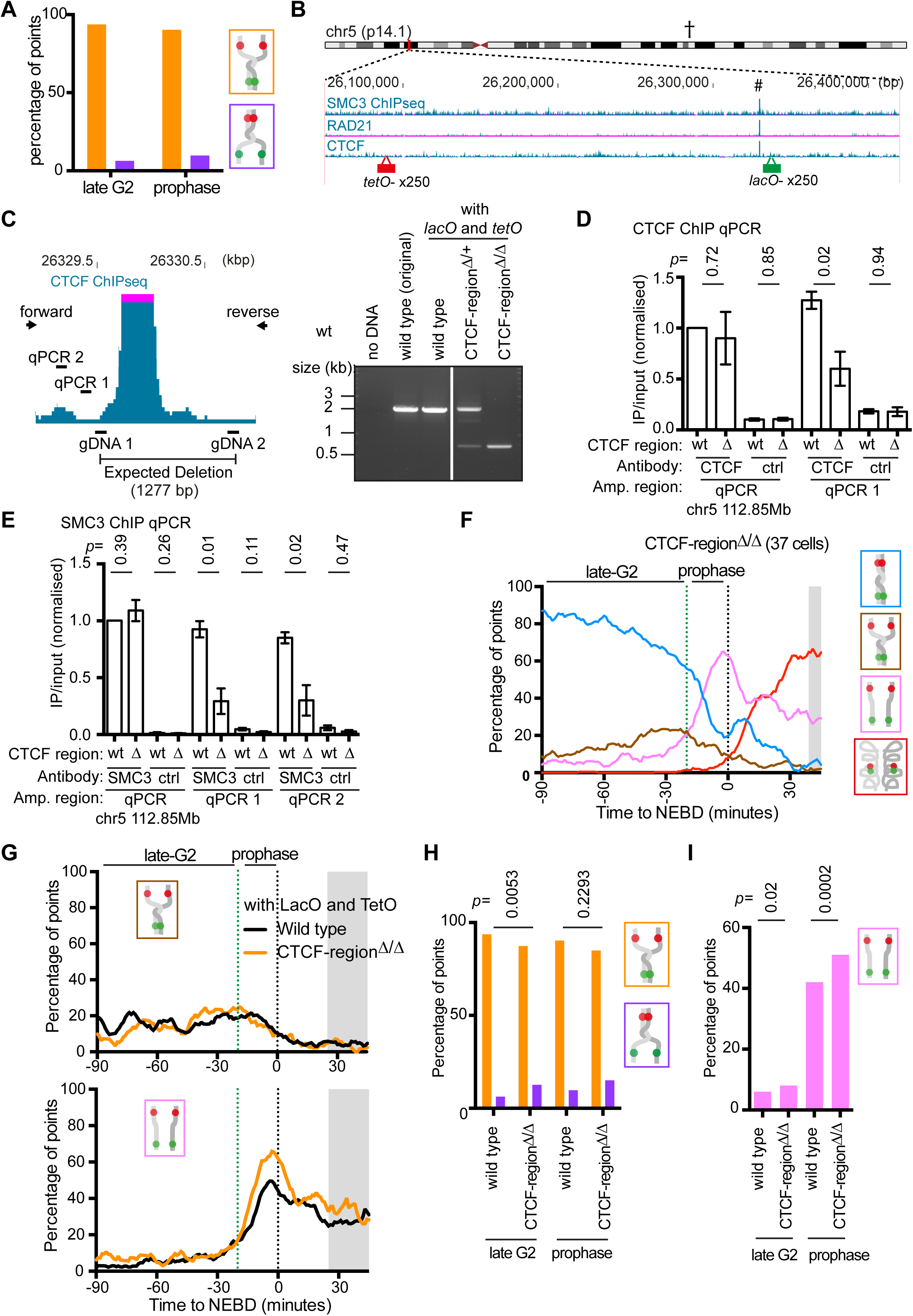
Local reduction of a cohesin peak on a chromosome leads to precocious sister chromatid resolution during prophase. (A) Graph shows the proportion of *tet* operator (orange colour) and *lac* operator (purple colour) sister separation amongst all “partial resolution” configurations in wild-type cells (TT75) during late G2-phase or prophase. The number of analysed points were 381 and 175 for late G2 or prophase, respectively. (B) The ChIP-seq (chromatin immunoprecipitation followed by high throughput DNA sequencing) data show the distribution of SMC3, RAD21, and CTCF along the genomic region where *lac* and *tet* operator arrays were inserted. The ChIP-seq data are taken from published data (Consortium, 2012) and their GEO accession numbers are GSM935384 (SMC3), GSM935571 (RAD21) and GSM733785 (CTCF). (C) The map (left) shows zoom-in of the chromosome region highlighted with # in (B), which contains a peak of SMC3, RAD21 and CTCF in ChIP-seq. The CTCF peak in ChIP-seq is shown in this map (purple highlights the data out of the y-axis scale). The map also shows positions of i) DNA sequences (gDNA1 and gDNA2), against which single guide RNA was designed for CRISPR-Cas9 genome deletion, ii) “forward” and “reverse” PCR primers that were designed to check the genome deletion (see DNA electrophoresis, right), and iii) the genome intervals (qPCR1 and qPCR2) that were amplified following ChIP in (D) and (E). In the DNA electrophoresis (right), wild-type cells showed a single band of an approximately 2kb PCR product while homozygous deletion (TT108 cells) showed a single band of approximately 0.7kb. A heterozygote deletion showed both bands. TT108 was generated by deleting the CTCF-binding sites in TT75 cells (see Figure 1(B)). (D) and (E) Graph shows results of ChIP followed by quantitative PCR (ChIP-qPCR). An antibody against CTCF or SMC3, or a non-specific antibody, was used for ChIP with wild-type (wt) or CTCF-region^Δ/Δ^ (Δ) cells (TT75 and TT108 respectively). The genome intervals (qPCR1 and qPCR2 in (C)) were amplified by PCR following ChIP. A region on chromosome 5 (112.85Mbp) indicated by † in (B) and seen Figure S3(D), where CTCF and cohesins are enriched, was also amplified by PCR as a control. For each sample, the yield (immunoprecipitated[IP]/input) was calculated and normalised to the yield at the control region in wild-type cells. The experiment was carried out three times and the mean and standard error of the mean for each condition is shown. *p* values for difference between wt and ∆ were obtained by unpaired *t*-tests. (F) The proportion of each configuration of the fluorescent reporter for CTCF-region^Δ/Δ^ cells (TT108) was plotted over time. TT108 cells were arrested by a double thymidine block, released from the block and observed by live-cell microscopy, as in Figure 1(F). Colour coding is indicated on the right-hand side. Data from individual cells are shown in Figure S3 (E). The start of prophase and NEBD are indicated as in Figure 2(E). The grey shaded area and data smoothing are as in Figure 1(F). (G) The proportion of the brown “partially resolved” (top) or pink “resolved” (bottom) states for wild-type or CTCF-region^Δ/Δ^ cells. These data were extracted from the data sets shown in Figure 1(F) and 3(F). The end of G2-phase and the start of prophase are indicated by the green dotted line and NEBD by the black dotted line. The grey shaded area is as Figure 2(F). (H) Graph shows the proportion of *tet* operator (orange colour) and *lac* operator (purple colour) sister separation amongst all “partial resolution” states in wild-type and CTCF-region^Δ/Δ^ cells during late G2 phase or prophase. *p* values were obtained using the *chi*-square statistical test, and the number of analysed points were 381 and 317 for wild-type and CTCF-region^Δ/Δ^ cells during late G2 phase, respectively and 175 and 93 for wild-type and CTCF-region^Δ/Δ^ cells during prophase, respectively. (I) The proportion of the pink “resolved” state for wild-type or CTCF-region^Δ/Δ^ cells during late G2-phase or prophase as indicated. These data were extracted from the data shown in Figure 1(F) and 3(F). *p* values were obtained using the *chi*-square statistical test, and the number of analysed points were 2406 and 2035 for wild-type and CTCF-region^Δ/Δ^ cells during late G2 phase, respectively and 991 and 680 for wild-type and CTCF-region^Δ/Δ^ cells during prophase, respectively. See also Figure S3.

To investigate the outcome of a reduced cohesin level at this region, we deleted a 1277 bp DNA sequence corresponding to the CTCF enrichment site containing three CTCF-binding consensus sites (Ziebarth et al., 2013). For this, we used a CRISPR-Cas9 deletion technique with two guide RNA sequences; one on each side of the CTCF binding sites (Figure 3C, left). Deletion of this region was confirmed by PCR and DNA sequencing on two homologous chromosomes of a selected clone (Figure 3C, right; Figure S3C), and was designated as “CTCF-region^∆/∆^”. ChIP with antibodies against CTCF or SMC3 was followed by quantitative PCR (qPCR), which amplified the vicinity of the deletion region (Figure 3C, left and S3D). This confirmed that deletion of the CTCF binding sites resulted in reduction of chromosome-bound CTCF and SMC3 by 53 and 67%, respectively, in this region (Figure 3D and E).

In the CTCF-region^∆/∆^ strain, we scored how fluorescent dots changed their configuration from late G2 to prometaphase, as we did in Figure 1E and F (Figure 3F and S3E). In the CTCF-region^∆/∆^ cell line, the overall fraction of the brown “partially-resolved” state was similar to wild-type control (Figure 3G, top), but there was a slight increase of sister separation of the *lac* operator array (green fluorescent dot), in late G2 phase (Figure 3H). Moreover, the CTCF-region^∆/∆^ cell line showed an earlier and greater increase of the pink “resolved” state during prophase, compared with wild-type (Figure 3G, bottom; Figure 3I). We conclude that a local reduction of the cohesin level leads to precocious sister chromatid resolution during prophase. As far as we know, this result provides the first evidence for a local effect of cohesin enrichment on sister chromatid cohesion in non-centromeric regions, i.e. enriched cohesins at a chromosome site locally contribute to robust cohesion. Moreover, while it was known that cohesins at CTCF binding sites promote chromatin loops in interphase (Dixon et al., 2012; Rao et al., 2014), our result suggests that cohesins at CTCF binding sites also contribute to robust sister chromatid cohesion. The precocious sister chromatid resolution happened mainly during late G2 in RAD21-depleted cells, and during prophase in CTCF-region^∆/∆^ cells (compare Figure 2F and 3G). This difference is probably due to remaining cohesins in the vicinity of the *lac* operator array in the CTCF-region^∆/∆^ cell line (Figure 3B and E) and normal levels of cohesins at other chromosome sites in this cell line.

### Sister chromatid resolution is established in prophase and maintained during prometaphase, relying on topoisomerase II activity

During DNA replication, chromosomes incur high levels of supercoiling that results in catenated sister chromatids. Therefore, in addition to cohesin removal, topoisomerase II (topo II), which catalytically removes catenanes from DNA, is required for sister chromatid resolution (Piskadlo and Oliveira, 2017; Pommier et al., 2016). Using our assay system, we next studied how chromosome re-organisation in early mitosis is affected by the specific catalytic inhibitor of topo II, ICRF-193 (Ishida et al., 1991). By observing metaphase chromosomes, we first confirmed the effect of ICRF-193 on our cells by analysing chromosome morphology. Following ICRF-193 treatment, the chromosomes looked tangled in the majority of metaphase cells (Figure 4A) suggesting that sister chromatid resolution was indeed defective, after topo II inhibition, as previously reported (Ishida et al., 1991).

**Figure 4:**
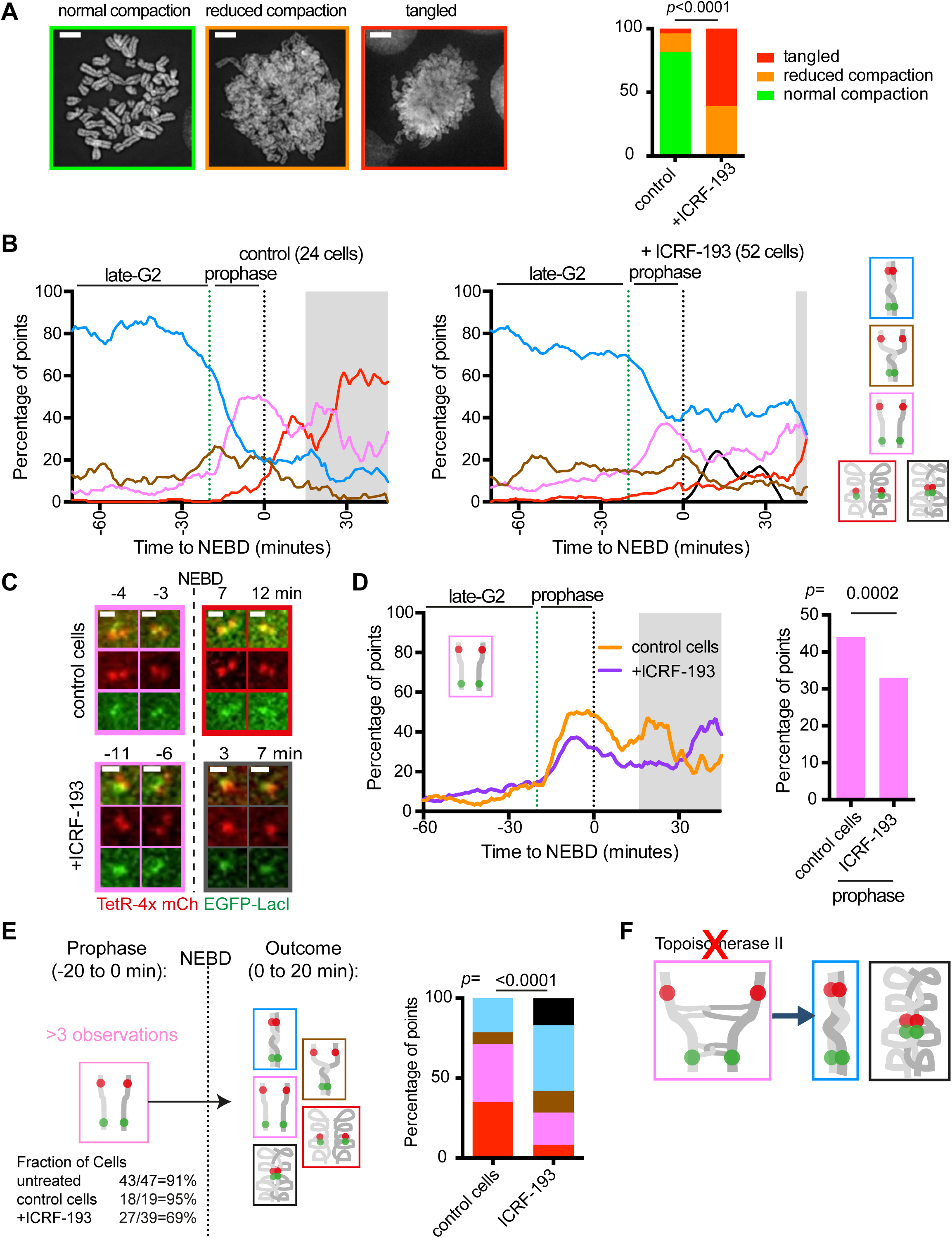
Topoisomerase II is required to stabilise resolved sister chromatids during prometaphase. (A) Metaphase spread analysis of ICRF-193 (topo II inhibitor)-treated and control cells. The images of representative metaphase spreads are shown on the left with the frame indicating the colour coding. The scale bar represents 5 µm. The proportion of each class of spread is shown on the right. The *p* value was obtained by the *chi*-square test and the number of spreads analysed were 27 and 23 for control and ICRF-193 treated cells, respectively. (B) The proportion of each configuration of the fluorescent reporter in control cells and cells treated with ICRF-193. TT75 cells were arrested by a double thymidine block, released from the block, treated with MK1775 and ICRF-193 (or only with MK1775 for control) as shown in Figure S4(A), and observed by live-cell microscopy as in Figure 1(F). Colour coding is shown on the right. Data from individual cells is shown in Figure S4(B). The grey shaded area and data smoothing are as in Figure 1(F). (C) Representative live-cell images of the fluorescent reporter in a control or ICRF-193 treated cell. The frame colours of the images indicate the configuration of the reporter evaluated according to Figure 1(D). Times indicated are relative to NEBD. (D) The proportion of the pink “resolved” state for control or ICRF-193 treated cells plotted against time (left) or during prophase (right). These data were extracted from the data sets shown in Figure 4(B). On the left, the end of G2-phase, the start of prophase and NEBD are indicated as in Figure 2(F). The grey shaded area is as in Figure 2(F). On the right, the *p* value was obtained by *chi*-square test, and the number of analysed time points were 366 and 961 for control and ICRF-193 treated cells, respectively. (E) The change in the fluorescent reporter configuration following NEBD was scored. Diagram (left) shows the pipeline for assessing individual cells; only those where the pink “resolved” state was observed for at least 4 time points during prophase (the fraction of such cells is indicated at the bottom) were assessed further. The proportion of each state during the 20 minutes following NEBD for the selected cells is shown for control or ICRF-193 treated cells. *p* value was obtained by *chi*-squared test and the number of analysed time points were 142 and 254 for control and ICRF-193 treated cells, respectively. (F) Diagram explaining the outcome with the inhibited topoisomerase II activity. With reduced topo II activity, DNA catenation may still remain after sister chromatid resolution is complete overall, which may subsequently de-stabilise largely resolved sisters, leading to reversion to the “unresolved” configuration. See also Figure S4.

We then compared the change in configuration of fluorescent dots from late G2 to early mitosis in the presence and absence of ICRF-193 (Figure 4B, C; Figure S4A, B). Since ICRF-193 treatment leads to engagement of the G2/M checkpoint (Downes et al., 1994; Ishida et al., 1994), we bypassed the checkpoint by using the WEE1 inhibitor, MK-1775 (Hirai et al., 2009) (Figure S4A). MK-1775 treatment itself (control) did not significantly affect the change in configuration of fluorescent dots (Figure 4B, left and S4B, left; compare with Figure 1E and F). In contrast, treatment with both ICRF-193 and MK-1775 (simply designated ICRF-193 treatment below) caused mild reduction of the pink “resolved” state during prophase (Figure 4D). ICRF-193 treatment also led to co-localisation of all four fluorescent dots after NEBD (black in Figure 4B, right), which we interpret as an abnormal “non-resolved and compacted” state (Figure S4C). Moreover, along the time course of individual ICRF-193 treated cells, the pink “resolved” state often reverted to the blue (and black) “non-resolved” state after NEBD (Figure 4C, bottom; Figure 4E; Figure S4B, right). When topo II activity is reduced, DNA catenation may still remain after overall completion of sister chromatid resolution in this region, and may subsequently de-stabilise largely resolved sisters, leading to such reversion (Figure 4F). With ICRF-193 treatment, chromosome compaction (the red “resolved and compacted” state plus the black “non-resolved and compacted” state) could proceed after NEBD, but not as efficiently as in control, since some cells showed reversion to the blue state as mentioned above.

We reproduced this phenotype of ICRF-193 treatment without using the WEE1 inhibitor; i.e. we arrested cells in G2/M with a CDK1 inhibitor, released them from the arrest (which bypassed G2/M checkpoint) and added ICRF-193 (Figure S4D), and this gave similar results to Figure 4B–E (Figure S4E, F). Overall we find that topo II activity is required to resolve sister chromatids in prophase, a role also observed by others (Gimenezabian et al., 1995; Liang et al., 2015; Nagasaka et al., 2016). In addition we have found a novel role of topo II in stabilising and maintaining resolved sister chromatids during prometaphase.

### Distinct roles of condensin I and II in sister chromatid resolution and chromosome compaction

Condensin I and II protein complexes (Figure 5A) play distinct roles in the re-organisation of mitotic chromosomes (Hirano, 2012). Next, we investigated the roles of condensin I and II using our assay system of real-time live cell imaging. In order to deplete cells of either condensin I or condensin II, we used siRNAs against NCAPD2 or NCAPD3 respectively. Depletion of these proteins was confirmed by western blotting (Figure 5B). We characterised the configuration of the fluorescent dots in these cells over time (Figure S5A and B) and plotted proportions of each configuration against time relative to NEBD (Figure 5C). NCAPD2-depleted cells showed an increase of the pink “resolved” state during prophase, with a similar timing to cells treated with a control siRNA (Figure 2F and Figure 5D, left). In contrast, NCAPD3-depleted cells showed a delay in the increase of the pink state, compared with control cells (Figure 5D, left). Intriguingly, both NCAPD2-depleted and NCAPD3-depleted cells showed a delay in the increase of red “compacted” state, relative to control cells; the extent of this delay was similar in NCAPD2-depleted and NCAPD3-depleted cells (Figure 5D, right).

**Figure 5:**
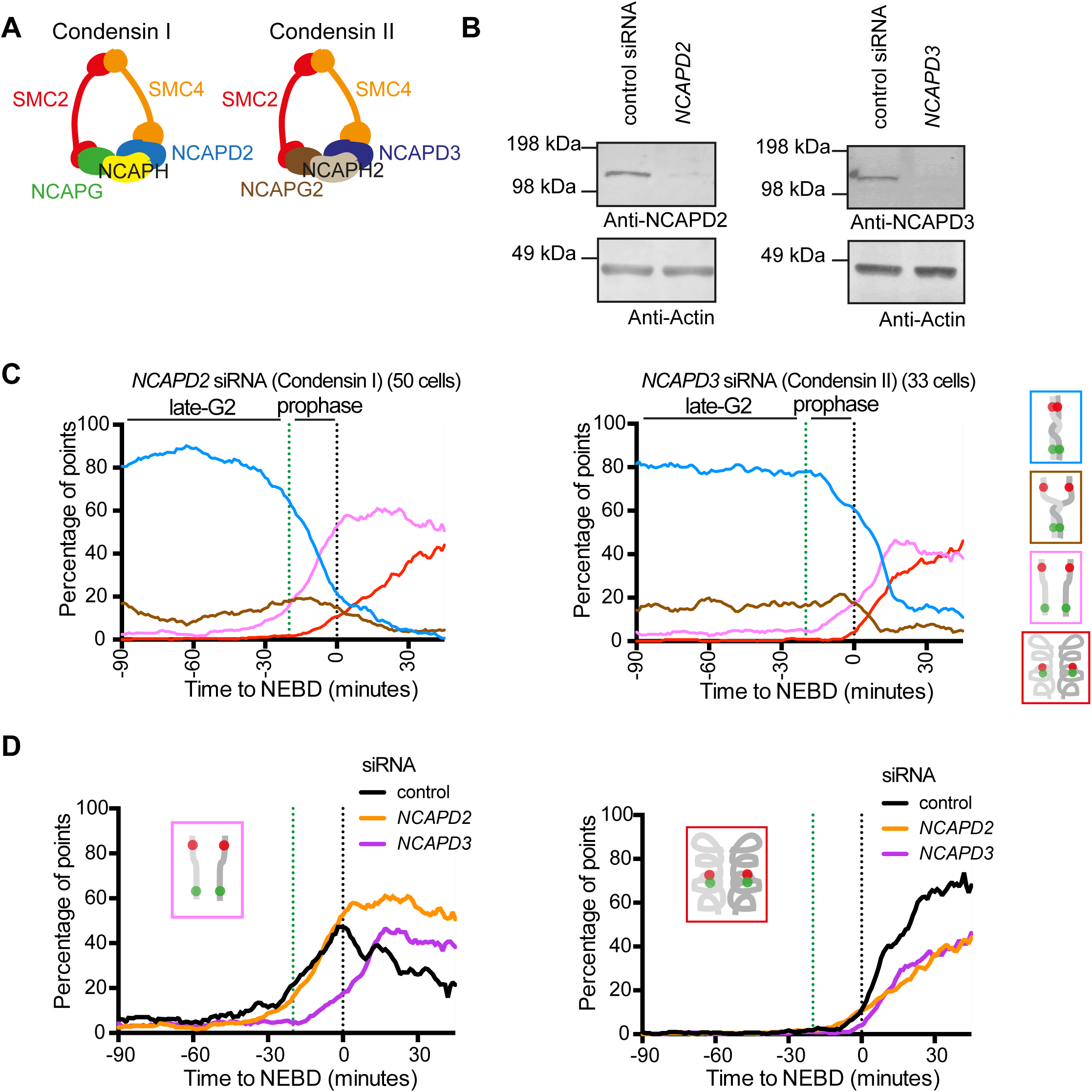
Condensin I and II play distinct roles in sister chromatid resolution and compaction. (A) Diagram showing composition of condensin I and II. Both contain the subunits SMC2 and SMC4 and are distinguished by the unique subunits NCAPG/H/D2 (condensin I) or NCAPG2/H2/D3 (condensin II). (B) Western blotting analysis of NCAPD2 (condensin I; left) or NCAPD3 (condensin II; right) proteins following treatment with the indicated siRNA for 48 hours. Re-probing with an actin antibody shows a loading control. (C) The proportion of each configuration of the fluorescent reporter for cells depleted for NCAPD2 or NCAPD3. TT75 cells were treated with NCAPD2 or NCAPD3 siRNA for 48 hrs, arrested by a double thymidine block, released from the block and observed by live-cell microscopy as in Figure 2(E). Colour coding is shown on the right. Data from individual cells is shown in Figure S5(A) and (B). Late G2 phase, prophase and NEBD are indicated as in Figure 1(F). Data smoothing are as in Figure 1(F). (D) The proportion of the pink “resolved” (left) or red “compacted” (right) state with control, NCAPD2 or NCAPD3 siRNA. These data were extracted from the data sets shown in Figure 2(E) and 5(C). Late G2 phase, prophase and NEBD are indicated as in Figure 2(F). See also Figure S5.

The results suggest that NCAPD3 (condensin II)-depleted cells show a defect in sister chromatid resolution. In contrast, NCAPD2 (condensin I)-depleted cells showed no delay in sister chromatid resolution, but did show a delay in chromosome compaction. Thus we conclude that condensin II and I play distinct roles in sister chromatid resolution and chromosome compaction. However, the exact extent of condensin II and I in promoting sister chromatid resolution and chromosome compaction, or the relative contribution of condensin II and I in facilitating these processes are still unknown. To address these issues, we need to understand the kinetics of these processes more quantitatively.

### Quantitative analyses reveal the specific timing of chromatid resolution and chromosome compaction

To analyse kinetics of sister chromatid resolution and chromosome compaction more quantitatively, we developed a stochastic model that describes the following two transitions in configuration of fluorescent dots; a) from the blue “non-resolved” state (including the brown “partially resolved” state) to the pink “resolved” state, and b) from the pink to the red “compacted” state (Figure 6A). We define that the transitions are stochastic with constant rates r_1_ (blue-to-pink) and r_2_ (pink-to-red) and constrained in time by the start time (ST) before NEBD and the time delay (TD) after ST, respectively (Figure 6A and B). A larger ST value means sister chromatid resolution occurs earlier, while a larger TD value means chromosome compaction happens with a larger delay after sister chromatid resolution (Figure 6B). The model proportions of blue, pink and red patterns were computed over 10,000 simulations for a given set of parameters. We found the best-fitting parameters by minimising the mean-square difference between the model and the data (calculated between −50 and +30 min, relative to NEBD) in each experimental condition (Figure 6C). Uncertainties of the best-fitting parameters were estimated by bootstrapping data, with the median reported as the best parameter estimate (Figure 6D, E). More details about this mathematical modelling are found in Experimental Procedures.

**Figure 6:**
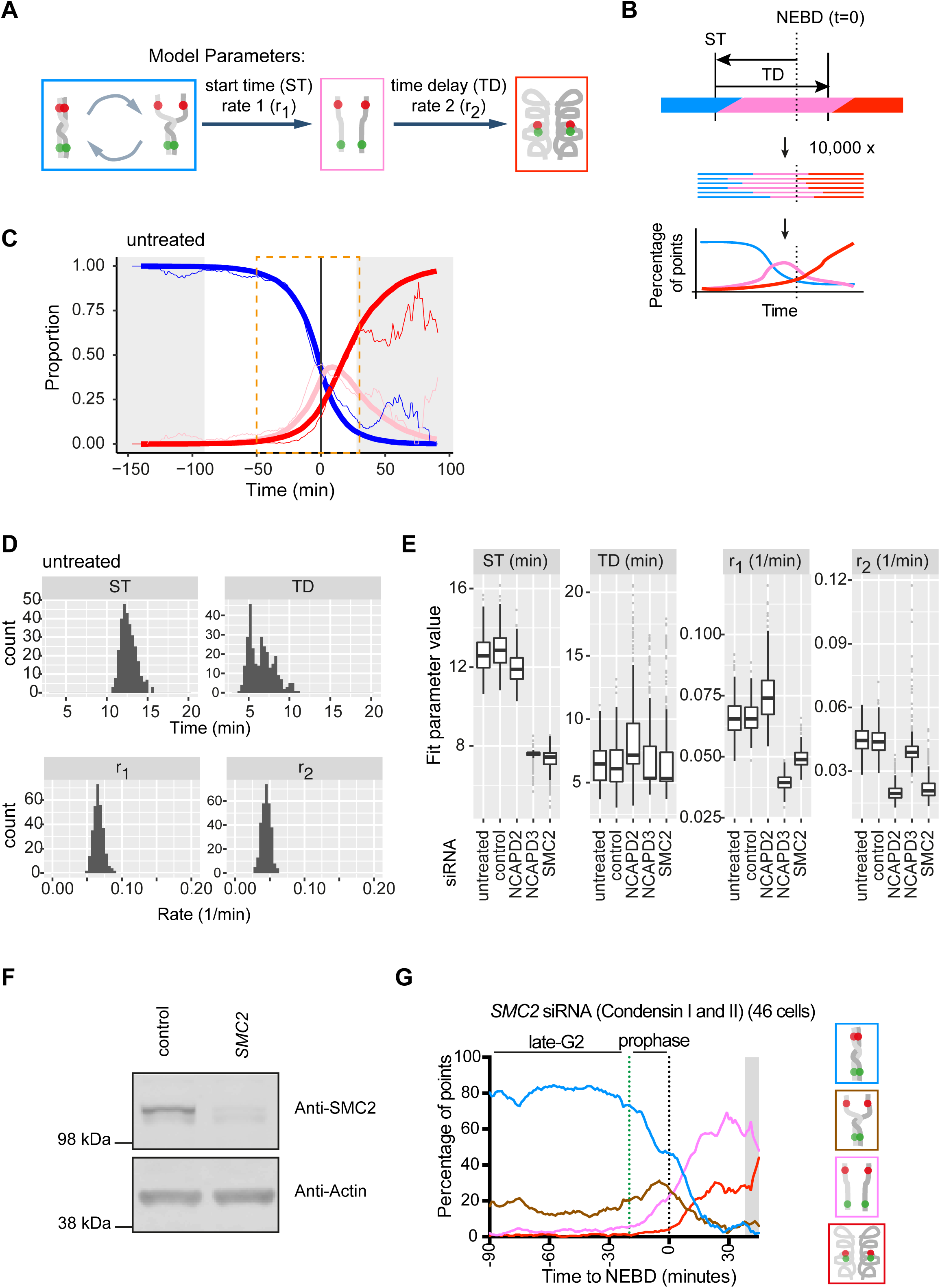
Mathematical modelling of sister chromosome resolution and compaction. (A) Diagram showing the main features of the model and the parameters that determine the transition between the different states. (B) Schematic showing how the model is calculated. For a given set of parameters (time constraints ST, TD and transition rates r_1_ and r_2_) 10,000 simulated cell timelines are created. Each timeline consists of a sequence of configurations (marked as blue, pink or red) as a function of time. The outcome of the model is the proportion of each configuration at each time point. The colour codes describing configurations are as in (A). (C) Example of untreated data (thin curves) with the best-fitting model (thick curves). The curves represent the proportion of each configuration in the fluorescent reporter system at each time point. The data curves were smoothed by a running mean with window size of 5 minutes for presentation only (that is, fitting was performed on unaltered data). The colour codes of lines are as in (A). The grey shaded area is as in Figure 1(F). The orange dotted box indicates the region over which the mean-square difference between the data and the model was calculated. (D) Example of bootstrap results to assess the uncertainty of fit parameters. In each bootstrap data were sampled with replacement and the best fit was found. 300 bootstraps were performed to create the distributions shown. See more details in Experimental procedures. (E) Box and whisker plots of fit parameters for the indicated conditions, obtained from bootstrapping (see D). The median (thick horizontal line) represents the best estimate of the parameter. (F) Western blotting analysis of SMC2 (condensin I and II) protein following treatment with SMC2 or control siRNA for 48 hours. Re-probing with an actin antibody shows a loading control. (G) The proportion of each configuration of the fluorescent reporter for cells depleted for SMC2. TT75 cells were treated with SMC2 siRNA for 48 hrs, arrested by a double thymidine block, released from the block and observed by live-cell microscopy as in Figure 2(E). Colour coding is shown on the right. Data from individual cells is shown in Figure S6(C). Late G2 phase, prophase and NEBD are indicated as in Figure 1(F). Data smoothing and grey shaded area are as in Figure 1(F). See also Figure S6.

As expected, the best-fitting parameter values obtained for untreated and control siRNA conditions were very similar (Figure 6E; S6A, B). In these ‘wild-type’ cells, values of ST and TD were 12–13 min and 6–7 min, respectively, suggesting that the final stage of sister chromatid resolution (transition to the pink “resolved” state) starts 12–13 min before NEBD, and chromosome compaction (transition to the red “compacted” state) begins 6–7 min later. In NCAPD3-depleted cells, ST was reduced by 5–6 min and r_1_ was reduced by about 40% relative to the control, which suggests a delay (by 5–6 min) and inefficiency (down to about 60%) in sister chromatid resolution (Figure 6E; S6A, B). On the other hand, NCAPD3-depleted cells showed an almost normal value of TD. This means the time interval between sister chromatid resolution and chromosome compaction remains largely the same even if the resolution is delayed (and occurs inefficiently), implying that resolution is a pre-requisite of compaction. Nonetheless, once chromosome compaction starts in NCAPD3-depleted cells, it proceeds with almost normal kinetics since r_2_ is almost unchanged.

In contrast, NCAPD2-depleted cells showed similar values of ST and r_1_ to control values, indicating largely normal sister chromatid resolution (Figure 6E; S6A, B). In NCAPD2-depleted cells, TD was also very similar to control values but r_2_ was reduced by about 55%, suggesting no significant delay but considerable inefficiency (down to about 45%) in chromosome compaction. Thus our quantitative analyses suggest that condensin II (NCAPD3) and condensin I (NCAPD2) specifically promote sister chromatid resolution and chromosome compaction, respectively, and our modelling reveals how much efficiency or time is lost when these factors are depleted.

To validate the above mathematical model, we made the following prediction: If we deplete SMC2, which is a common subunit of condensin I and II (Figure 5A), we would see the combined phenotypes of NCAPD2-and NCAPD3-depleted cells, i.e. the combined changes in parameter values. To test this prediction, we depleted SMC2 using siRNA which was confirmed by western blotting (Figure 6F). We then analysed the configuration of fluorescent dots by live-cell imaging of the SMC2-depleted cells (Figure 6G and S6C), applied the above mathematical model, and fitted parameter values for ST, TD, r_1_ and r_2_ (Figure 6E and S6A). In SMC2-depleted cells, the values of ST and r_1_ were similar to those in NCAPD3-depleted cells while the value of r_2_ was similar to that in NCAPD2-depleted cells. In other words, when either NCAPD2-or NCAPD3-depleted cells showed changes in parameter values, SMC2-depleted cells also showed very similar changes in these parameter values (Figure 6E). This supports that our mathematical model indeed provides accurate parameter values to represent timing and efficiency of sister chromatid resolution and chromosome compaction.

## Discussion

Human cells undergo dramatic changes in chromosome organisation in early mitosis, i.e. sister chromatids are resolved and chromosomes become compacted. To address timing, dynamics and potential coordination of these two events, we developed a novel real-time assay using live-cell imaging. Compared with other assays for mitotic chromosome reorganisation, our assay has the following three major advantages: First, our assay allows analyses of chromosome configuration with high spatial resolution – we visualise two neighbouring chromosome sites (100–250 kbp interval) as small fluorescent dots, and investigate the change in their configuration over time. Observation with high spatial resolution allows analyses of dynamic local chromosome changes, which are often obscured by using more standard observation of global chromosome changes. Second, our assay allows us to analyse changes of chromosome configuration with high temporal resolution – we acquire live-cell images every minute and align time course data of individual cells relative to NEBD. By observing events in individual cells with high temporal resolution, we can analyse rapidly changing events. Third, our method allows analyses of both sister chromatid resolution and chromosome compaction in one assay. By analysing both together, we can address their relative timing and potential coordination. Other assays usually focus on only one of the two; for example, Hi-C analyses enables detailed genome-wide study of chromosome compaction, but provides little information about sister chromatid resolution. Taking these advantages of our assay, we found novel regulation and dynamics of mitotic chromosome organisation – we highlight and discuss the following three findings:

First, we found that sister chromatid resolution already starts in late G2 phase and completes during prophase. Intriguingly, sister chromatids repetitively cycle through unresolved and partially resolved states before completing their resolution. The cyclical behaviour we observe might reflect multiple attempts to remove the stably bound cohesin from chromosomes. It could also reflect the motion of a dynamic population of cohesin moving locally along chromosomes (Busslinger et al., 2017; Ocampo-Hafalla et al., 2016).

Second, we found that a local reduction of cohesin accumulation by deletion of CTCF binding sites led to precocious sister chromatid resolution. This suggests that local accumulation of cohesins along chromosome arms correlates with the robustness of sister chromatid cohesion, a finding that previous global studies could not uncover. Moreover, although it was known that cohesins enriched at CTCF binding sites promote intra-chromatid looping during interphase (Dixon et al., 2012; Rao et al., 2014), it was unknown if they also support sister chromatid cohesion. Our results provide the first evidence for this notion. Cohesins at CTCF binding sites may switch their roles between intra-chromatin looping and sister chromatid cohesion. It will be intriguing to study whether such switching indeed occurs and, if so, how it is regulated.

Third, by analysing both sister chromatid resolution and chromosome compaction in one assay, we successfully quantified relative timing and regulation of the two events. On average, the resolution begins 12–13 min prior to NEBD and the compaction follows 6–7 min later. The resolution seems to be a pre-requisite of condensation since a delay in resolution leads to a similar amount of delay in compaction. Nonetheless, efficiency of the resolution and compaction is under independent regulation by condensin II and I, respectively. Recent computer modelling suggests that chromatin looping could drive not only chromosome compaction, but also sister chromatid resolution (Goloborodko et al., 2016). If so, defects in the resolution with condensin II depletion, but not with condensin I depletion, could be explained by the observation that condensin II promotes chromatin looping earlier than does condensin I (Gibcus et al., 2018). Meanwhile, defects in the compaction with condensin I depletion, but not with condensin II depletion, could be explained by the result that condensin II and I promote large and small chromatin looping, leading to chromosome axial shortening and chromosome compaction, respectively (Gibcus et al., 2018). Overall our results provide an important framework for temporal and molecular regulation of sister chromatid resolution and chromosome compaction.

To visualise selected chromosome loci, we used *tet* and *lac* operator arrays, inserted at targeted chromosome loci. Alternative methods for labelling selected chromosome loci in live cells use the nuclease-deficient Cas9 (dCas9) and single guide RNAs (sgRNA) (Chen et al., 2013; Ma et al., 2015). While the operator-based method requires establishment of stable cell clones carrying operators at targeted loci, the dCas9/sgRNA method could work with transient transfection. Thus, by using different sgRNAs, the dCas9/sgRNA method could allow us to analyse more chromosome regions in a short space of time. However, this method has been mainly used to visualise repetitive DNA sequences that provide workable signal intensities. Since our assay relies on visualisation of two neighbouring chromosomal loci (100 to 250kb apart), and suitably spaced unique repetitive sequences are not available, currently such dCas9/sgRNA methods would not be suitable to carry out the experiments we described. Nonetheless, the sensitivity of dCas9/sgRNA method is improving rapidly (Maass et al., 2018; Neguembor et al., 2018) and may allow more routine visualisation of non-repetitive sequences in the near future. Analyses of a larger number of chromosome regions in various cells and in various conditions, with live-cell imaging, would broaden our knowledge about mitotic chromosome organisation.

## Acknowledgements

We thank J. Swedlow, T. Owen-Hughes, W.C. Earnshaw, T. Hirota, W. Bickmore, J. Chubb and members of Tanaka, Hochegger and Fukagawa labs for discussion; T. Hori, D. Sherratt, K. Nasmyth, R.Y. Tsien, M. Davidson for reagents; G. Ball, P. Appleton, S. Swift, M. Tsenkov, D. Alexander and S. Ejidiran for technical help; and G. Barton for supervising the Data Analysis Group. This work was supported by an ERC advanced grant (322682), the Wellcome Trust (096535 and 097945), Cancer Research UK (C28206/A14499) and the Medical Research Council (K015869). T. U. T is a Wellcome Trust Principal Research Fellow. H. H. is a Cancer Research UK Senior Research Fellow.

## Experimental Procedures

### Cell Culture

The human cell line HT-1080 (obtained from ATCC) and derivative cell lines were cultured at 37°C and 5.0% CO_2_ under humidified conditions in DMEM (with L-glutamine), 10% fetal bovine serum, 100 U/ml penicillin, and 100 mg/ml streptomycin. Media and supplements were obtained from Invitrogen. For microscopy the above medium was replaced with Fluorobrite DMEM medium (Invitrogen) supplemented with 10% fetal bovine serum, 2mM L-glutamine, 1mM pyruvate and 25mM HEPES.

The DT40 cell line BM-lacO-20K-19 (Hori et al., 2013) and derivative cell lines were cultured at 37°C and 5.0% CO_2_ under humidified conditions in RPMI 1640 (with L-glutamine), 10% fetal bovine serum, 1% chicken serum, 100 U/ml penicillin, and 100 mg/ml streptomycin. Media and supplements were obtained from either Invitrogen or Lonza. For microscopy the above medium was replaced with colourless RPMI-1640 (phenol red-free) medium supplemented with 10% fetal bovine serum, 1% chicken serum, 2mM L-glutamine, 1mM pyruvate and 25mM HEPES.

Transfection of plasmids into HT-1080 and derivative cell lines was facilitated with Fugene HD according to manufacturers instructions (Promega). Cells were transfected in single wells of a 6-well dish using 4.5µl Fugene HD and 1.5µg plasmid (3:1 ratio). Selection was performed 24 to 48 hours after transfection using puromycin (Sigma; 0.3 µg/ml), blasticidin (Invivogen; 2 µg/ml), hygromycin (Roche; 60 µg/ml), histidinol (Sigma; 2 mg/ml), and G418 (Sigma; 300 µg/ml).

Transfection of DT40 cells was performed by electroporation of 1.0 × 10^7^ cells with 10–15 mg linearized plasmid using 0.4 cm Gene Pulsar cuvettes and Gene Pulsar electroporation apparatus (Bio-Rad) at 550 V and 25 mF. Selection was performed using puromycin (final concentration 0.5µg/ml) or histidinol (final concentration 1mg/ml).

Protein knockdown by siRNA was carried out using Lipofectamine (Invitrogen) according to manufacturers instructions. In all cases 0.01nmol of siRNA with 6µl of lipofectamine and 200µl of Optimem (Invitrogen) was added to cells in 2 ml of medium in 6-well or 3cm microscopy dishes. The medium containing the siRNA was replaced and fresh siRNA added every 24 hours. Cells for analysis by western blotting or microscopy were used between 48 and 60 hours after first addition of siRNA. siRNAs were obtained from Ambion or Eurofins. The sequences were as follows; WAPL (5’-CGG ACU ACC CUU AGC ACA AdTdT-3’); RAD21 (5’-AUA CCU UCU UGC AGA CUG UdTdT-3’); NCAPD2 (5’-CGU AAG AUG CUU GAC AAU UTT-3’); NCAPD3 (no.1 5’-GAU AAA UCA GAG UAU CGU ATT-3’ or no. 2 5’-GAA CAG CGA UUC AAC AUC ATT-3’); SMC2 (no. 1 5′-GAA UUA GAC CAC AAC AUC AdTdT-3′ or no. 2 5’-CUA UCA CUC UGG ACC UGG AdTdT-3’); non-specific (5’-UAA CGA CGC GAC GAC GUA ATT-3’).

Synchronisation of human cells at the G1/S-phase boundary was achieved using a double-thymidine block (Figure S2D). Essentially, 0.1 × 10^6^ cells were seeded in 2 ml of medium in 6-well dishes or 3cm glass-bottomed microscopy dishes (WPI) 16 to 24 hours before the treatment. Thymidine was then added at a final concentration of 2.5mM and incubated for 16 hours. Thymidine was then removed and cells were washed 3 × 2 ml with fresh medium. Cells were incubated for a further 8 hours before 2.5mM Thymidine was added. Cells were then incubated for 12 – 16 hours and Thymidine was then removed and cells were washed 3 × 2 ml with fresh medium and incubated for a further 5 to 10 hours to observe cells in late S-phase or G2/M-phase as appropriate.

Synchronisation of human cells at the G2/M-phase boundary was achieved using the CDK1-inhibitor RO-3306 (Millipore; Figure S4D). Essentially, 0.1 × 10^6^ cells were seeded in 2 ml of medium in 6-well dishes or 3cm glass-bottomed microscopy dishes (WPI) 16 to 24 hours before the treatment. RO-3306 was then added at a final concentration of 9 µM and cells were incubated for 12 hours. RO-3306 was then removed and cells were washed with 4 × 2ml PBS before 2ml fresh medium was added.

The Topoisomerase II inhibitor ICRF-193 (Sigma) was used at a concentration of 2µg/ml and the WEE1 inhibitor MK1775 (Selleckchem) was used at a concentration of 0.5µM. The DNA stain SiR-DNA (tebu-bio) was used at a concentration of 200nM and was added to cells up to 12 hours before imaging.

### Plasmids

For integrating *tet* operator arrays into human chromosome 5 the plasmids pT2770 and pT2707 were used. pT2707 contains the Cas9 gene and the specific guide DNA (sequence: ACG GGT TCT TGT CCG TCC CA) which was cloned into pGeneArt-CRISPR-Nuclease according to manufacturer instructions (Invitrogen; A1175). In this case, and all other cases mentioned, the Zhang lab resources at http://www.genome-engineering.org/crispr/ were used to aid gRNA design. pT2770 was created to target the *tet* operator arrays to the region and contained homology to genomic regions upstream and downstream of the selected gDNA site of pT2707. The homology arms were amplified from genomic DNA by PCR using primer pairs Chr5-26Mb-5’F (AAA AAA CTC GAG CGA TCG TGT CTT GGG GAT GTT CCA CGG GTA C) Chr5-26Mb-5’R (AAA AAA AGA TCT TTG GAT CCA CGG ACA AGA ACC CGT CTT CAG CTG) and Chr5-26Mb-3’F (AAA AAA GGA TCC TTA GAT CTC CCA CGG CCA TGA AAA TGT GGG CTC) Chr5-26Mb-3’R (AAA AAA GCG GCC GCC TTT CTT GAC ACA TTG TTG GGA ACC) and sequentially cloned into ploxPuro (Arakawa et al., 2001). *tet* operator arrays (250 repeats) with non-repetitive 10bp DNA sequences between each repeat from pLAU44 (Lau et al., 2003) and the puromycin resistance gene (from ploxPuro) were then sequentially cloned in between the regions of homology to create pT2770. At each stage restriction digestion and sequencing were used to confirm cloning.

For integrating *lac* operator arrays into human chromosome 5, 250kb upstream of the *tet* operator arrays, the plasmids pT2846 and pT2837 were used. pT2837 contains the Cas9 gene and the specific guide DNA (sequence: CAT TTA GGT TTT TCA CGT AC) was cloned into pGeneArt-CRISPR-Nuclease according to manufacturer instructions (Invitrogen; A1175). pT2846 was created to target the *lac* operator arrays to the region and contained homology to genomic regions upstream and downstream of the selected gDNA site of pT2837. The homology arms were amplified from genomic DNA by PCR using primer pairs 5Chr5-26Mbii-F (AAA AAC TCG AGA TGC TAA GTG TGG GAG GGC AAT TTC) 5Chr5-26Mbii-R (AAA AAG GAT CCT TGT CGA CGT ACT GGG ATA ATA GGA ACA TTT GAA AC) and 3Chr5-26Mbii-F (AAA AAA GGA TCC TTA GAT CTG TGA AAA ACC TAA ATG ACA CCA TCA CC) 3Chr5-26Mbii-R (AAA AAA GCG GCC GCC TGC CTC TCT CTC TCA TAC ACA TGT G) and sequentially cloned into ploxBlast (Arakawa et al., 2001). *lac* operator arrays (250 repeats) with non-repetitive 10bp DNA sequences between each repeat from pLAU43 (Lau et al., 2003) and the blasticidin resistance gene (from ploxBlast) were then sequentially cloned in between the regions of homology to create pT2846. At each stage restriction digestion and sequencing were used to confirm cloning.

For integrating *lac* operator arrays into human chromosome 5, 750kb upstream of the *tet* operator arrays, the plasmids pT2838 and pT2847 were used. pT2838 contains the Cas9 gene and the specific guide DNA (sequence: TAG GCT TCA CCG TAG TAT CT) was cloned into pGeneArt-CRISPR-Nuclease according to manufacturer instructions (Invitrogen; A1175). pT2847 was created to target the *lac* operator arrays to the region and contained homology to genomic regions upstream and downstream of the selected gDNA site of pT2838. The homology arms were amplified from genomic DNA by PCR using primer pairs 5Chr5-26Mb-Fiii (AAA AAC TCG AGA TTA ACT TCC ACT ACT CTA CTA GAG CTG) 5Chr5-26Mb-Riii (AAA AAG GAT CCT TGT CGA CAT CTT GGA TAC TAC CTA CGT ATG TAT G) and 3Chr5-26Mb-Fiii (AAA AAA GGA TCC TTA GAT CTA CTA CGG TGA AGC CTA CAT AGA C) 3Chr5-26Mb-Riii (AAA AAA GCG GCC GCC ACT GTA TTA TTT TCC TAG AGC TGC CC) and sequentially cloned into ploxBlast (Arakawa et al., 2001). *lac* operator arrays (250 repeats) with non-repetitive 10bp DNA sequences between each repeat from pLAU43 (Lau et al., 2003) and the blasticidin resistance gene (from ploxBlast) were then sequentially cloned in between the regions of homology to create pT2847. At each stage restriction digestion and sequencing were used to confirm cloning.

For integrating *tet* operator arrays into the chicken (DT40) Z chromosome the plasmid pHH100TetO was used. pHH100TetO contains homology to genomic regions upstream and downstream of the selected target site. The homology arms were amplified from genomic DNA by PCR using primer pairs 100TetLA5Not (TAT AGC GGC CGC CCT CAG ATT GTT CAA ACA TTA ATG AGA TGC) 100TetLA3Bgl (ATA AGA TCT GGA TCC CCA TAT CTG AAA TCC AAA TGT TTA CAA AAT) and 100TetRA5Bgl (TAT AAG ATC TAC AAC CTA TTG AGC AGT TGA AGG TGG AAG G) 100TetRA3Xho (TAT ACT CGA GGC TAG TGC TGC TGG ATT ATC CAG AAG CTC C) and sequentially cloned into pBLUESCRIPT. Next, *tet* operator arrays from pLAU44 (Lau et al., 2003) and the puromycin resistance gene were sequentially cloned in between the regions of homology to create pHH100TetO.

For visualising *tet* operator arrays in live human or chicken cells a plasmid expressing TetR-4mCherry from the beta-actin promoter was created. For this, *tetR* (Michaelis et al., 1997) and 4 copies of mCherry (Renshaw et al., 2010) were cloned into pExpress (Arakawa et al., 2001) along with the histidinol resistance gene from pJE59 (Eykelenboom et al., 2013). Restriction digestion and sequencing was used to confirm cloning.

For visualising *lac* operator arrays in live human or chicken cells the plasmid pEGFP-lacI-NLS was used (Hori et al., 2013). For visualising replication factories in live human cells the plasmid pmCerulean-PCNA-19-SV40NLS-4 was used and was a gift from the Davidson lab (Addgene plasmid 55386).

For deleting the specific CTCF-binding site in human cells (detailed in the main text) the plasmids pT3093 and p3099 were used. pT3093 and pT3099 contain the Cas9 gene and the specific guide DNAs (sequences: GAC TTA GTC CCT ACC TCA CA and AAT CAC TGT GAG CCT GCC TA) which were cloned separately into pGeneArt-CRISPR-Nuclease according to manufacturers instructions (Invitrogen; A1175). pT3093 and pT3099 were designed to cleave upstream and downstream of the targeted CTCF-binding site respectively.

### Cell lines

The human HT-1080 derived cell line containing *tet* operator and *lac* operator arrays, separated by 250kbp of DNA, and expressing TetR-4mCherry and EGFP-LacI was designated TT75 and was created as follows. HT-1080 cells were transfected with pT2707 and pT2770 and puromycin resistant clones were obtained. Targeting of the *tet* operator arrays to the desired genomic location was confirmed using the primers 5F (CTT GTG ACA TGA CCT TCT AAA TAG AGT GC), 5R (CAC TGC ATT CTA GTT GTG GTT TGT CC), 5ctrlF (AAA AAA CTC GAG CGA TCG TGT CTT GGG GAT GTT CCA CGG GTA C), 5ctrlR (AAA AAA AGA TCT TTG GAT CCA CGG ACA AGA ACC CGT CTT CAG CTG). This cell line was designated TT51. This cell line was then sequentially transfected with pT2415 and pEGFP-lacI-NLS with selection for histidinol and hygromycin resistance respectively. Finally, these cells were transfected with pT2837 and pT2846 and blasticidin resistant clones were obtained. Targeting of the *lac* operator arrays to the desired genomic location was confirmed using the primers 5Fii (GCA GGT GCA TGG GAA TAC AAG TGT TG), 5Rii (CTC ATC AAT GTA TCT TAT CAT GTC TGG ATC), 5ctrlF (AAA AAC TCG AGA TGC TAA GTG TGG GAG GGC AAT TTC) and 5ctrlR (AAA AAG GAT CCT TGT CGA CGT ACT GGG ATA ATA GGA ACA TTT GAA AC).

The human HT-1080 derived cell line containing *tet* operator and *lac* operator arrays, separated by 750kbp of DNA, and expressing TetR-4mCherry and EGFP-LacI was designated TT68 and was created as follows. TT51 (containing *tet* operator arrays and whose construction is described in the previous paragraph) was sequentially transfected with pT2415 and pEGFP-lacI-NLS with selection for histidinol and hygromycin resistance respectively. These cells were then transfected with pT2838 and pT2847 and blasticidin resistant clones were obtained. Targeting of the *lac* operator arrays to the desired genomic location was confirmed using the primers 5Fiii (CTC ATT ATC TGT ACA TTT CTT TGC ATC G), 5Riii (CTC ATC AAT GTA TCT TAT CAT GTC TGG ATC), 5ctrlFiii (AAA AAC TCG AGA TTA ACT TCC ACT ACT CTA CTA GAG CTG) and 5ctrlRiii (AAA AAG GAT CCT TGT CGA CAT CTT GGA TAC TAC CTA CGT ATG TAT G) using a similar strategy to that shown in Figure S1B for integration of the *lac* operator array at an alternative location.

The TT75 derivative expressing Cerulean-PCNA was designated TT104 and was created by transfecting TT75 cells with pmCerulean-PCNA-19-SV40NLS-4 with selection for G418 resistance.

The TT75 derivative with a deletion of the CTCF-binding region close to the *lac* operator array was designated TT108 and was created by transfecting cells with pT3093 and pT3099. Stable clones were obtained and screened for the deletion by PCR using the primer pair CTCF-F (TGC ATT TTA AGT GCT CAC TAG AGG) and CTCF-R (GTG CCA TTC AGA ACA TTT TTA GAG). The region around the deletion was sequenced using the primer CTCF-R.

The DT40 cell line containing *lac* operator arrays and *tet* operator arrays was designated TT56 and was a derivative of BM-lacO-20K-19. BM-lacO-20K-19 was a DT40 cell line in which a *lac* operator array (512 repeats) had been targeted to a position 3.8 Mbp along the Z chromosome (Hori et al., 2013)(Figure S1D). The TT56 cell line was created by transfecting these cells with the plasmid pHHTetO100 and selecting for puromycin resistance. Stable clones were obtained and screened by Southern blotting; genomic DNA was digested with EcoRI and the probe was generated using the primer pair 100South-F TTT GCA GAG GTC CAT GGC TCC CCA ACC CAG 100South-R GTT AGC AAG CCT GCA ATA TCA AGA AAG GAG. A successfully targeted clone was then transfected with pT2415 and stable clones obtained by selecting for histidinol resistance.

### SDS PAGE and Western blotting

For western analysis, total cell extracts were obtained from cells grown in 6-well dishes and lysed in 30 – 60 µl of lysis buffer (20mM HEPES pH7.6; 400mM NaCl; 1mM EDTA; 25% glycerol; 0.1% NP-40) containing protease inhibitors (cOmplete EDTA-free; Roche). Lysates were quantified using Bradford reagent (Thermo; 1863028) and 30 µg of total protein for each sample was ran on pre-cast Bis-Tris 4–12% gradient gels (Invitrogen) and protein transferred to PVDF membrane (Amersham). Membranes were blocked in PBS containing 5% milk and were incubated with antibodies in PBS containing 2% BSA and 0.05% (w/v) sodium azide. Primary antibodies were used as follows: WAPL (Abcam; ab70741) 1 in 5000; RAD21 (Millipore; 05-908) 1 in 2000; NACPD2 (Sigma; HPA036947) 1 in 1500; NCAPD3 (Bethyl; A300-604A) 1 in 3000; SMC2 (Abcam; ab10412) 1 in 5000; Actin (Sigma; A5441) 1 in 20,000. Secondary antibodies were used as follows: Donkey-anti-mouse-800CW (LI-COR; 926-32212) 1 in 10,000; Donkey-anti-rabbit-680RD (LI-COR; 926-68073) 1:10,000. Signal from the secondary antibody was detected using a LI-COR Odyssey CLx.

### Metaphase spreads

After appropriate treatment approximately 2 × 10^6^ cells were collected and incubated in 5 ml of hypotonic solution (75 mM KCl) for 10 min at 37ºC. The cells were then incubated in 5 ml of cold fixative (methanol/acetic acid; 3:1) for 20 min at 37ºC. This fixation step was repeated and cells finally resuspended in approximately 200 µl of fixative solution and stored at −20ºC. For spreading, 50 µl of fixed cells were dropped onto glass slides and air dried. The spread chromosomes were mounted in Prolong Gold Antifade containing DAPI (Invitrogen) before imaging.

### Chromatin Immunoprecipitation (ChIP)

Cells were plated and grown in 10 cm dishes until confluent (1 – 2 × 10^6^ cells). For each ChIP approximately 4 × 10^6^ cells were used (2 × 10cm dishes). Cells were crosslinked with 1.42% (v/v) formaldehye for 10 minutes and quenched with 125mM for a further 10 minutes. Cells were washed three times in 10ml PBS and then scraped from the dishes. Cells were resuspended in 2 × 300µl lysis buffer containing 1% SDS, 10mM EDTA, 50mM Tris pH8.1 and protease inhibitors (cOmplete EDTA-free Roche) and chromatin was fragmented (to approximately 0.5 to 1.5 kb) using a Biorupter (Diagenode) at medium setting with total sonication time of 10 minutes (20 cycles of 30 seconds on/off). Lysates were cleared by centrifugation for 10 minutes at high speed and 50µl saved to create the input sample. The remaining lysates were diluted 1:10 in ChIP dilution buffer containing 1% TritonX-100, 2mM EDTA, 150mM NaCl, 20mM Tris pH8.1, 0.1% Brij-35. The lysates were then incubated for 1 hour at 4°C with 60µl Protein G beads. Beads were then removed and these lysates were incubated with 2µg of antibody (see below) overnight at 4°C. 60µl of Dynabeads which had been incubated overnight in PBS containing 0.5% (w/v) BSA were then added to the lysate plus antibody mix and incubated for a further 4 hours. The supernatant was then discarded and the beads washed twice with 1 ml of each of the following; wash buffer I (0.1% SDS, 1% TritonX-100, 2mM EDTA, 20mM Tris pH8.1, 150mM NaCl), wash buffer II (0.1% SDS, 1% TritonX-100, 2mM EDTA, 20mM Tris pH8.1, 500mM NaCl), wash buffer III (0.25M LiCl, 1% NP-40, 1% sodium deoxycholate, 1mM EDTA, 10mM Tris pH8.1) and TE buffer (10mM Tris pH8.1, 1 mM EDTA). Antibody-Protein-DNA complexes were then eluted in 2 × 100µl of Elution buffer (1% SDS, 0.1M sodium bicarbonate). Cross-links were then reversed for these samples and the input sample by adding 8µl 5M NaCl and incubating at 65°C overnight. Next 10µl of RNaseA (10mg/ml) was added and incubated at 37°C for 30 minutes and then 4µl 0.5M EDTA, 8µl Tris pH8.1 and 10 µl Proteinase K (15mg/ml) were added and incubated at 45°C for 2 hours. Finally, DNA was purified using minElute DNA purification columns (QIAGEN) and eluted in 20µl of elution buffer.

### Quantitative PCR (qPCR)

Quantitative PCR was carried out using an Eppendorf LightCycler 96 and RotorGene SYBRGreen PCR mix (Qiagen; 1054596) according to manufacturers instructions. For each experiment technical replicates were prepared for each sample. Input chromatin was diluted 1:100. In order to compare the overall pull-down efficiency between cell lines positive-control primers were used CTCF-pos-F (CGG AGT ATC AAG GCA TCA GTA A) and CTCF-pos_R (GGA ATC GCA CAG TTG AGA ATA AG). To assess pull-down of our region of interest two sets of query primers were used CTCF-qu-F1 (GTC ATG TCT TCA GTG CAT GAT TT), CTCF-qu-R1 (GGT AGG GAC TAA GTC TGT TTC G) and CTCF-qu-F2 (AGT GTC ATT AGT GCT TCC TTC T), CTCF-qu-R2 (GAGAATGCTCTGGCCTCTTT). After PCR ΔCq values for each sample were obtained by Cq_input_ – Cq_pull-down_, from this we calculated each as a fraction of input (2^−ΔCq^) to measure pull-down efficiency. Finally, each value was normalised as a fraction of the pull-down achieved for the positive control primers on chromatin from the wild-type cells.

### Microscopy and Image analysis

Time-lapse images were collected at 37°C with 5% CO_2_ using a DeltaVision ELITE microscope (Applied Precision), an UPlanSApo 100x objective lens (Olympus; numerical aperture 1.40), softWoRx software (Applied Precision) and either a CoolSNAP HQ or EMCCD Cascade II camera (both Photometrics). For HT-1080 derivative cells we acquired 25 z-sections 0.75µm apart. For DT40 cells we acquired 25 z-sections 0.5µm apart. During live cell imaging Cerulean, EGFP and mCherry signals were discriminated using the dichroic CFP/YFP/mCherry (52-850470-000 from API). For imaging chromosomes in live cells SiR-DNA (emission 674nm) and EGFP signals were discriminated using the dichroic DAPI/FITC/TRITC/Cy5 (52-852111-001 from API).

Fixed cell images were collected using the same DeltaVision ELITE microscope. For metaphase spreads 15 sections 0.2µm apart were acquired DAPI signal was detected using the dichroic DAPI/FITC/TRITC/Cy5 (52-852111-001 from API).

In all cases, after acquisition images were deconvolved using softWoRx software with enhanced ratio and 10 iterations. Analysis of individual cells was carried out using Imaris software (Bitplane). Statistical analyses were carried out using Graphpad Prism 6.0 software.

The configuration of the fluorescent *tet* and *lac* operator arrays in live cells was determined through time and in three-dimensions using Imaris imaging software (Bitplane). For both blue “non-resolved” and brown “partially resolved” states at least one operator (*tet* or *lac*) was observed as a single fluorescent object. To distinguish blue “non-resolved” and brown “partially resolved” states we looked at the second operator region such that: (1) if it was observed as (i) a single fluorescent object or (ii) appeared as two fluorescent objects whose centres were separated by less than 1 µm we classified it as the blue “non-resolved” state; or (2) if it was observed as two fluorescent objects whose centres were separated by more than 1µm we classified it as the brown “partially resolved” state. For both pink “resolved” and red “non-resolved” states both of the operator regions appeared as two separate fluorescent objects (four objects in total; two for *lac* and two for *tet*). To distinguish pink “resolved” and red “non-resolved” states we looked at the colocalisation of the *lac/tet* paired fluorescent objects such that: (1) if one pair (or both) showed less than 50% colocalisation we classified them as the pink “partially resolved” state; or (2) if both pairs showed more than 50% colocalisation we classified them as the red “compacted” state. In some cases, following NEBD, all four fluorescent dots showed more than 50% colocalisation over at least 5 consecutive timepoints and we classified this as a black “non-resolved and compacted” state.

### Encode data sets

ENCODE datasets (ENCODE Project Consortium, 2012) were visualised using genome browser at UCSC (**http://genome.ucsc.edu/**) with the Human Feb. 2009 (GRCh37/hg19) Assembly (Kent et al., 2002). The ENCODE Genome expression omnibus (GEO) accession numbers were as follows: RAD21 GSM935571 (Snyder lab, Stanford); SMC3 GSM935384 (Snyder lab, Stanford); CTCF GSM733785 (Bernstein lab, Harvard).

### Mathematical modelling

We developed a mathematical model that describes both the transition of chromosomes (or chromosomal regions) to the resolved configuration and the subsequent transition to a compacted state (Figure 6A and B). The model is stochastic and based on Poisson processes. The first and second transition can occur only after the time *t*_1_ and *t*_2_, respectively. These two times are randomly drawn from exponential distributions. Specifically, *t*_1_ takes place before NEBD, *t*_1_ = *t_NEBD_* − *R_e_*(*τ*_1_) and *t*_2_ happens after *t*_1_, *t*_2_ = *t*_1_ + *R_e_*(*τ*_2_), where *R_e_*(*τ*) is a random variable with cumulative distribution 1 − *e^−t^*^/^*^τ^*.

After *t*_1_ the transition to the resolved state can occur with a constant rate *r*_1_. After *t*_2_ and only when the first transition happened, the transition to the compacted state can happen with a constant rate *r*_2_. Hence, the model is controlled by four parameters, time scales *τ*_1_, *τ*_2_ and transition rates *r*_1_ and *r*_2_. Based on the rules outlined above a timeline for a cell is calculated as a sequence of states on a constant time step. The calculation is repeated for 10,000 cells and proportions of each state for each time point are found.

We fit the model to experimental data using Broyden–Fletcher–Goldfarb–Shanno (BFGS) algorithm implemented in *optim* function in R package (version 3.4.3). We minimize the sum of squares of differences between the model and the data in a fixed time window from −50 to +30 min relative to *t_NEBD_*.

To assess the uncertainty of fit parameters we performed a bootstrap. For a given data set we sampled with replacement data points within the time window and found the best-fitting parameters for each sample. The procedure was repeated 300 times, giving us a distribution of values for each parameter. From this we found the median as the central representation of the parameter and 95% confidence limits. Parameters *τ*_1_ and *τ*_2_ can be understood as typical time scales at which state transitions are allowed. For example, for untreated cells we find *τ*_1_ = 18.2 min. The corresponding half-life time (*τ* ln 2 = 12.6 min) tells us that, on average, half of the cells will be able to enter the resolved state 12.6 min before NEDB. For simplicity of interpretation we converted *τ*_1_ and *τ*_2_ values to half-life values in our figures and report them as start time (ST) and time delay (TD), respectively. Similarly, the rates *r*_1_ and *r*_2_ can be interpreted in terms of half-lives (1ln2) of decay into a transformed state or as constant probabilities of transition in a single time step, 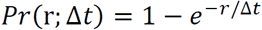. For example, *r*_1_ = 0.065 min^−1^ for untreated cells. This corresponds to a decay half-life of 10.7 min or the probability of transition in a single 1-min time step of 0.062.

The model was implemented in R. The software with documentation (R markdown document) are available from GitHub at https://github.com/bartongroup/MG_ChromCom.

---

**Figure S1.**
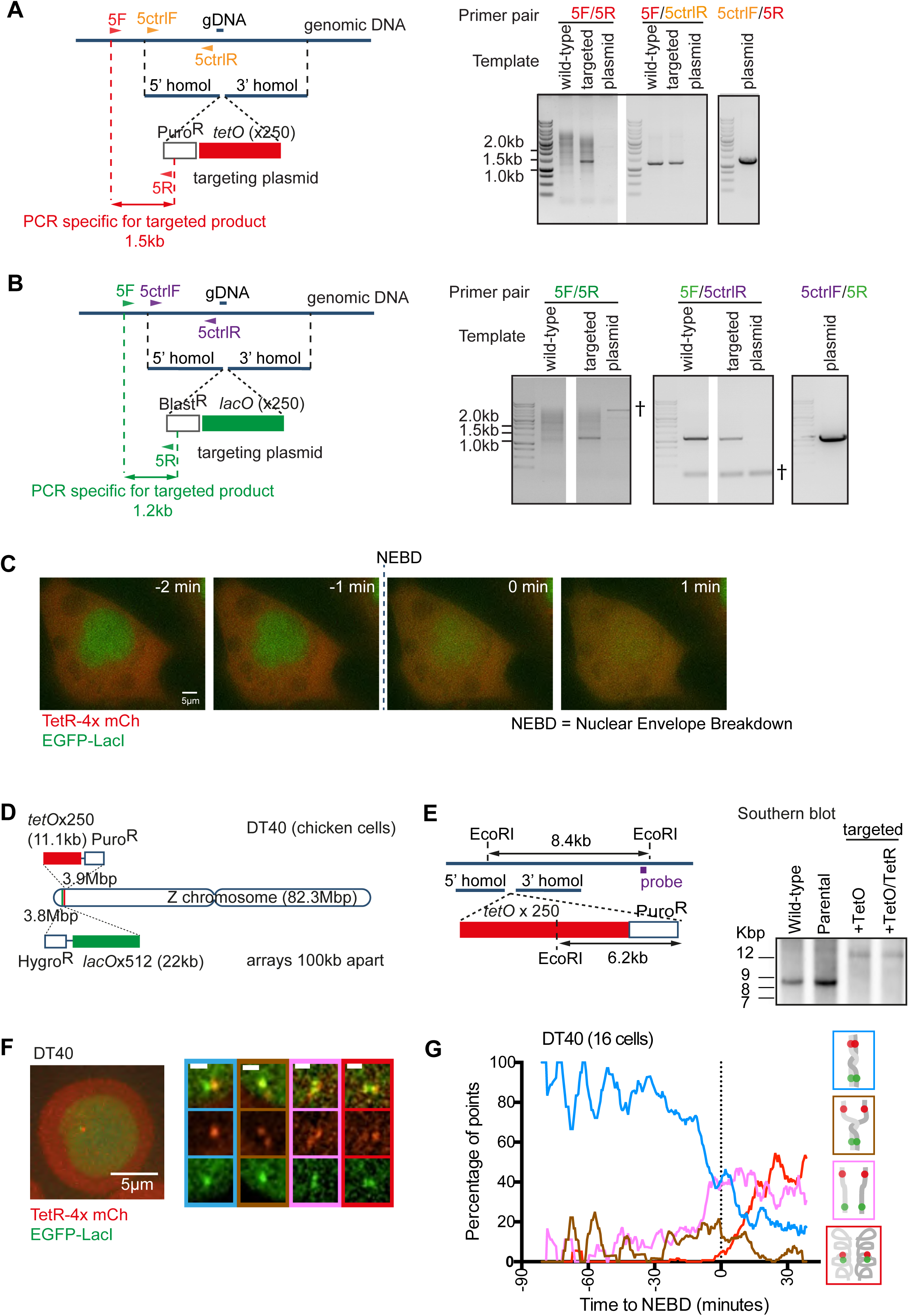
(related to Figure 1) (A) The map (left) shows the strategy for targeted integration of *tet* operator arrays to the region of chromosome 5 shown in Figure 1(A). The map also shows relative positions of (i) the target site of the guide RNA (gDNA), (ii) PCR primers 5F and 5R used to check for targeted integration (see DNA electrophoresis, right) and (iii) control PCR primers 5ctrlF and 5ctrlR for genomic or plasmid DNA (see DNA electrophoresis, right). The DNA electrophoresis (right) show the results of PCR using template DNA from the wild-type genome (wild-type), the genome with targeted integration (targeted), and, as control, a purified plasmid (plasmid) containing 5’ and 3’ homology sequences, puromycin resistance gene and *tet* operators (see map, left). The 5F/5R primer gives a single 1.5 kb product using ‘targeted’, but not ‘wild-type’ or ‘plasmid’ DNA as a template. In contrast, the control primer pair 5F/5ctrlR gives a 1.5 kb product using ‘targeted’ and ‘wild-type’ DNA as a template, but not using ‘plasmid’ DNA. Use of the control primer pair 5ctrlF/5R confirms that ‘plasmid’ DNA can give a PCR product (about 1.5 kb). Overall the PCR results demonstrate that, in selected cell clones, the construct with the *tet* operator array was correctly integrated as we designed. (B) The map (left) shows the strategy for targeted integration of *lac* operator arrays to the region of chromosome 5 shown in Figure 1(A). The map also shows relative positions of (i) the target site of the guide RNA (gDNA), (ii) PCR primers 5F and 5R used to check for targeted integration (see DNA electrophoresis, right) and (iii) control PCR primers 5ctrlF and 5ctrlR for genomic or plasmid DNA (see DNA electrophoresis, right). The DNA electrophoresis (right) show the results of PCR using template DNA from the wild-type genome (wild-type), the genome with targeted integration (targeted), and, as control, a purified plasmid (plasmid) containing 5’ and 3’ homology sequences, blasticidin resistance gene and *lac* operators (see map, left). The 5F/5R primer gives a single 1.2 kb product using ‘targeted’, but not ‘wild-type’ or ‘plasmid’ DNA as a template. In contrast, the control primer pair 5F/5ctrlR gives a 1.2 kb product using ‘targeted’ and ‘wild-type’ DNA as a template, but not using ‘plasmid’ DNA. Use of the control primer pair 5ctrlF/5R confirms that ‘plasmid’ DNA can give a PCR product (about 1.2 kb). Non-specific bands are indicated by †. Overall the PCR results demonstrate that, in selected cell clones, the construct with the *lac* operator array was correctly integrated as we designed. (C) Time lapse images of an individual cell undergoing NEBD. TT75 cells were arrested by a double thymidine block and subsequently released from the block. 8–10 hr after the release, images were taken every minute. Times indicated are relative to NEBD. Scale bar is 5µm. (D) Diagram depicting the location of *tet* and *lac* operator arrays introduced into the Z chromosome of DT40 cells using homologous recombination gene targeting methods. (E) The map (left) shows the targeting strategy for integration of *tet* operator arrays to the region of the Z chromosome shown in part (D). The map also shows relative positions of (i) EcoRI restriction sites in the genome (dark blue line, top) or the *tet* operator array region (red box, bottom), (ii) the position of the DNA probe (purple line) for the Southern blot screen. Southern blotting (right) using the indicated probe reveals a 12kb product for the targeted clones and a 8.4kb product for the wild-type cells. (F) Visualisation of these arrays by expression of TetR-4x mCherry and EGFP-LacI in DT40 cells (TT56) (left). Scale bar represents 5µm. Representative images of the major configurations of the fluorescent reporter observed in TT56 cells (right). Designated colour-codes for each are indicated in the image frames. Scale bar represents 1µm. (G) Behaviour of the fluorescent reporter observed in wild-type live-cells. Asynchronously growing TT56 cells were imaged every minute and, at each time point, the configuration of the reporter was determined and represented using the colour-codes depicted in (F). Data from individual cells was aligned according to the occurrence of nuclear envelope breakdown (NEBD; time zero) and proportions of each configuration at each time point was determined with smoothing applied by calculating the rolling mean proportion across 9 minutes. Colour coding for each configuration is indicated on the right-hand side.

**Figure S2:**
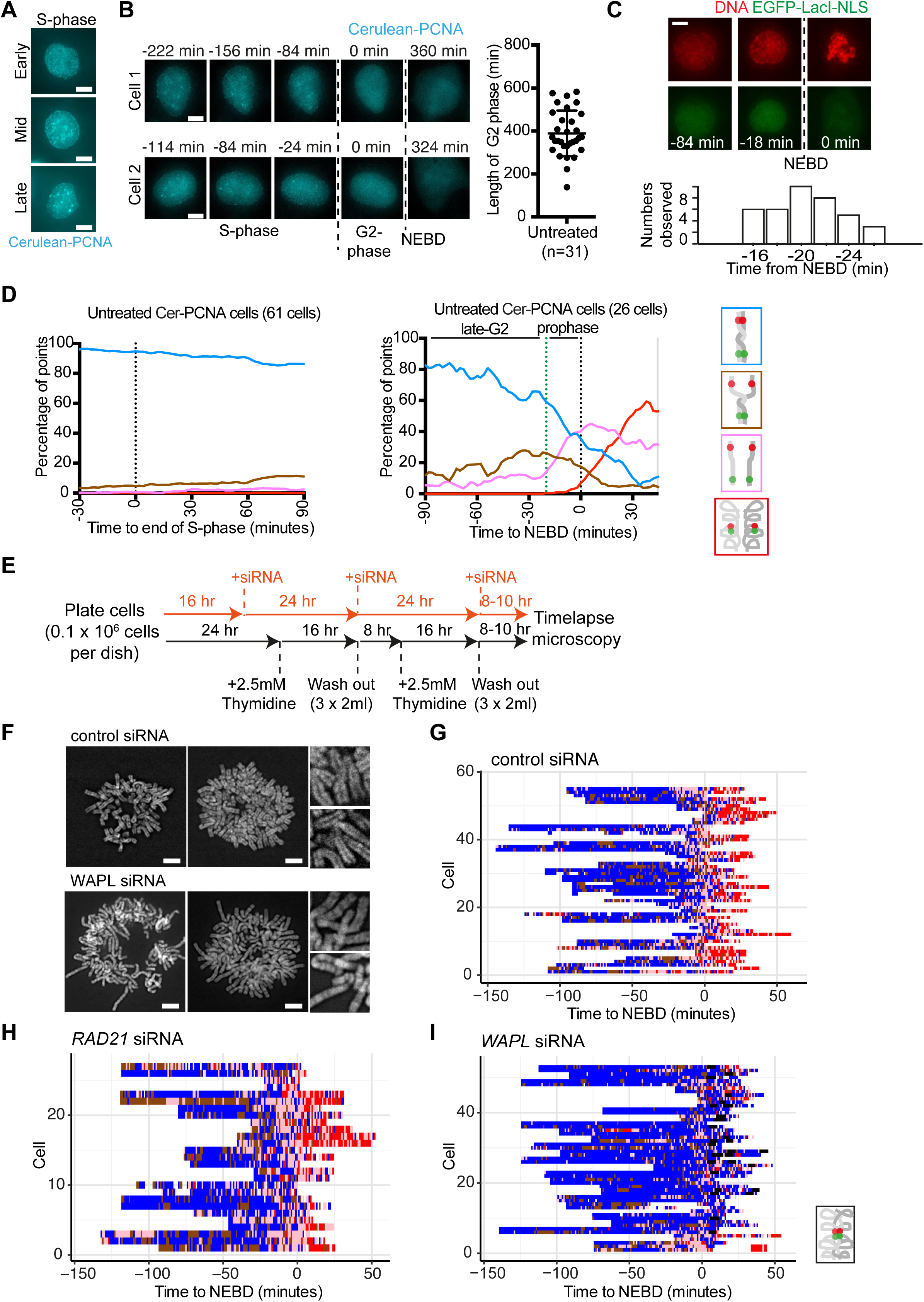
(related to Figure 2) (A) Snap shot images of Cerulean-PCNA observed in live asynchronously growing TT104 cells. Images show replicating cells with characteristic PCNA foci state of Early, Mid and Late S-phase. Scale bar represents 10µm. (B) Measuring the length of G2 phase in individual TT104 cells by observing Cerulean-PCNA behaviour. TT104 cells were arrested by a double thymidine block and subsequently released from the block. 5 hours after the release, images were taken every 6 minutes. On the left are time lapse images of Cerulean-PCNA times indicated are relative to the point of disappearance of the Cerulean PCNA foci (end of S-phase). NEBD was identified after the diffusion of free Cerulean PCNA throughout the cell. The phases of the cell cycle of the images are indicated. Scale bar represents 10µm. The G2-phase time period for individual cells (between the end of S-phase and NEBD) was measured and plotted (right). (C) Evaluating the length of prophase. TT75 cells were incubated with SirDNA and their images were acquired every 2 min. A representative cell is shown at the top. Timing of NEBD was defined as the time of dispersion of GFP-LacI signal from the nucleus to the whole cell (time zero). The start of prophase was evaluated (min relative to NEBD; graph on the bottom) as the time that condensed globular SirDNA signals were observed in >50% of the area of the nucleus, including at the nuclear periphery, in images projected to 2D. Before prophase, a small number of globular SirDNA signals were observed (see −84 min image) but these were unstable and disappeared within 4 min, in contrast to globular signals in prophase. From these results, we define that prophase starts at −20 min relative to NEBD. (D) Behaviour of the fluorescent reporter observed in wild-type live-cells. TT104 cells were synchronised as in part (B) and, 5 hours (late S and early G2) or 8–10 hours (late G2 and prophase) after the release images were taken every two minutes and, at each time point, the configuration of the reporter was determined and represented using the colour-codes depicted on right. Data from individual cells was aligned according to the end of S-phase (left plot) or according to NEBD (right plot) as determined according to part (B) and proportions of each configuration at each time point was determined with smoothing applied by calculating the rolling mean proportion across 9 timepoints. (E) Experimental procedure outline. siRNA was added to cells 16 hours after plating and then cells were synchronised in early S-phase using a double thymidine block. After each thymidine wash-out the relevant siRNA was added back to the cells as indicated. (F) Metaphase spread analysis of WAPL-depleted and control cells. Asynchronously growing TT75 cells were treated with WAPL or control siRNA for 48 hours before fixation and metaphase spread preparation. Shown are representative images obtained. Scale bar is 5µm. (G), (H) and (I) Behaviour of the fluorescent reporter as observed in individual live-cells, treated with control, RAD21 or WAPL siRNA (respectively), plotted across the Y-axis. TT75 cells were treated with siRNA, arrested by a double thymidine block and subsequently released from the block. 8–10 hr after the release, images were taken every minute and, at each time point, the configuration of the reporter was determined and represented using the colour-codes depicted in Figure 1(C) and (D). Data from individual cells was aligned as in Figure 1(E). White spaces represent time points where configuration could not be determined.

**Figure S3:**
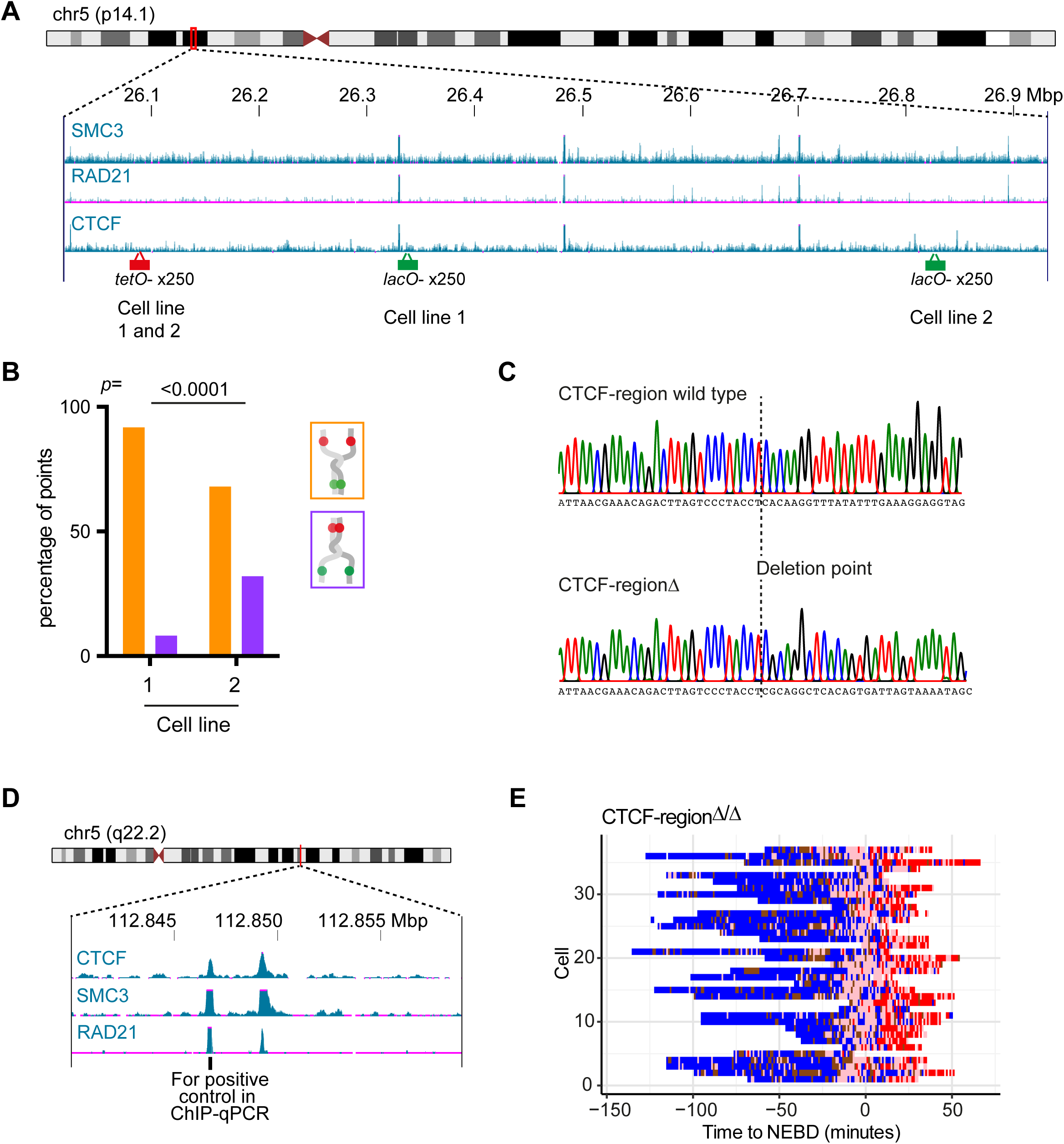
(related to Figure 3) (A) The ChIP-seq (chromatin immunoprecipitation followed by high throughput DNA sequencing) data show the distribution of SMC3, RAD21, and CTCF along the genomic region where *lac* and *tet* operator arrays were inserted in cell line 1 (TT75 and derivatives including TT104) and cell line 2 (TT68). The ChIP-seq data are taken from published data (Consortium, 2012) and their GEO accession numbers are GSM935384 (SMC3), GSM935571 (RAD21) and GSM733785 (CTCF). (B) Graph shows the proportion of *tet* operator (orange colour) and *lac* operator (purple colour) sister separation amongst all “partial resolution” configurations in cell line 1 (TT104) and cell line 2 (TT68) during late G2-phase and prophase. In each case microscopy images were taken by time-lapse every 2 minutes. The *p* value was obtained using the *chi*-square statistical test, the number of analysed time points were 208 and 50 for cell line 1 and cell line 2, respectively. (C) Sequencing chromatogram of the CTCF-binding region from wild-type (TT75) or CTCF-region^Δ/Δ^ (TT108) cells. Sequencing was carried out across the gDNA1 region as shown in Figure 3(C). The junction of the deletion is indicated by the dotted line. The CTCF-regionΔ shown here contains deletion of the 1277bp region between gDNA 1 and gDNA 2. (D) The ChIP-seq (chromatin immunoprecipitation followed by high throughput DNA sequencing) data show the distribution of SMC3, RAD21, and CTCF along the genomic region where the qPCR positive control primers are situated. The ChIP-seq data are taken from published data (ENCODE Project Consortium, 2012) and their GEO accession numbers are GSM733785 (CTCF), GSM935384 (SMC3) and GSM935571 (RAD21). (E) Behaviour of the fluorescent reporter as observed in individual CTCF-region^Δ/Δ^ live-cells, plotted across the Y-axis. TT108 cells were arrested by a double thymidine block and subsequently released from the block. 8–10 hr after the release, images were taken every minute and, at each time point, the configuration of the reporter was determined and represented using the colour-codes depicted in Figure 1(C) and (D). Data from individual cells was aligned according to Figure 1(E). White spaces represent time points where configuration could not be determined.

**Figure S4:**
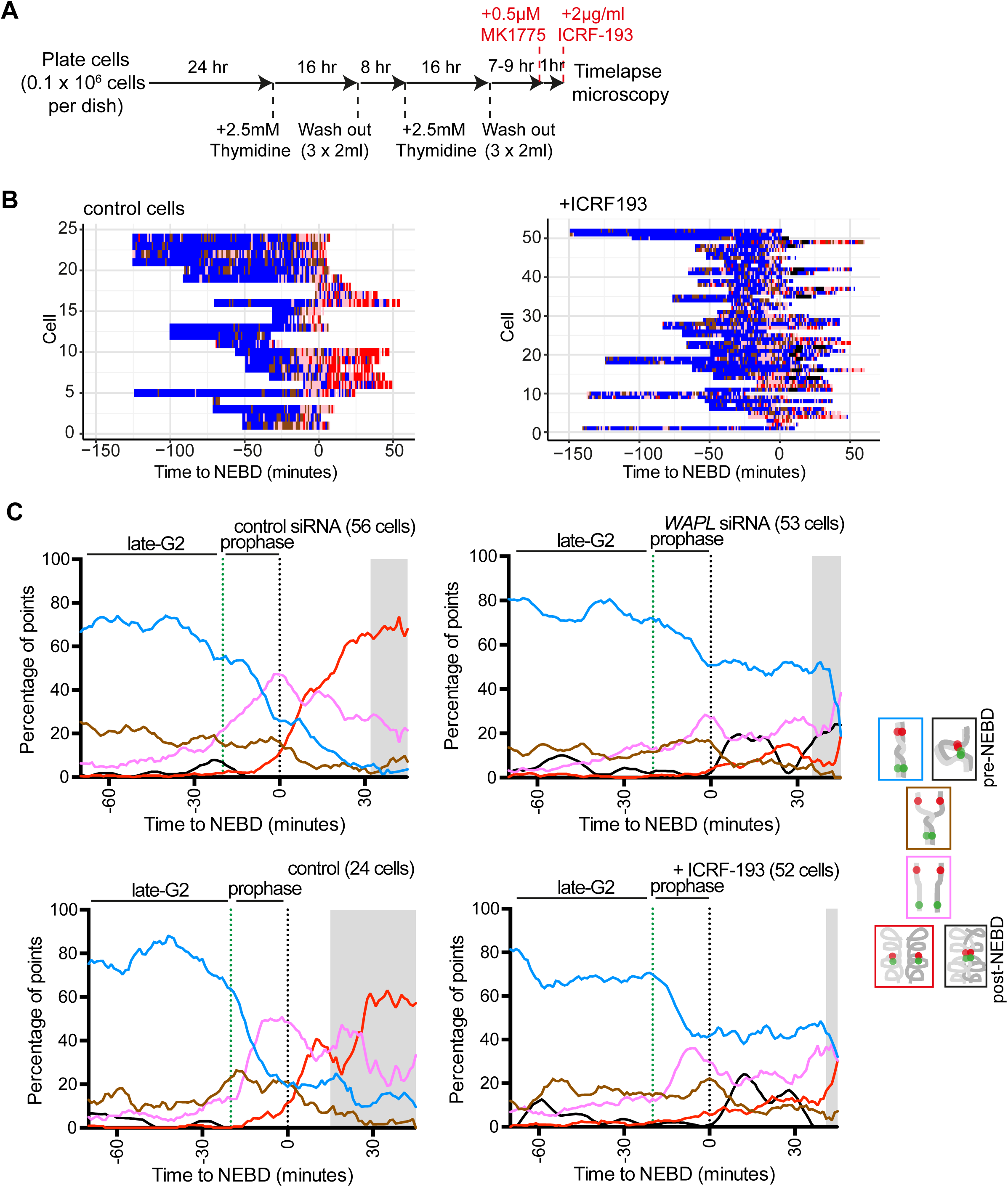
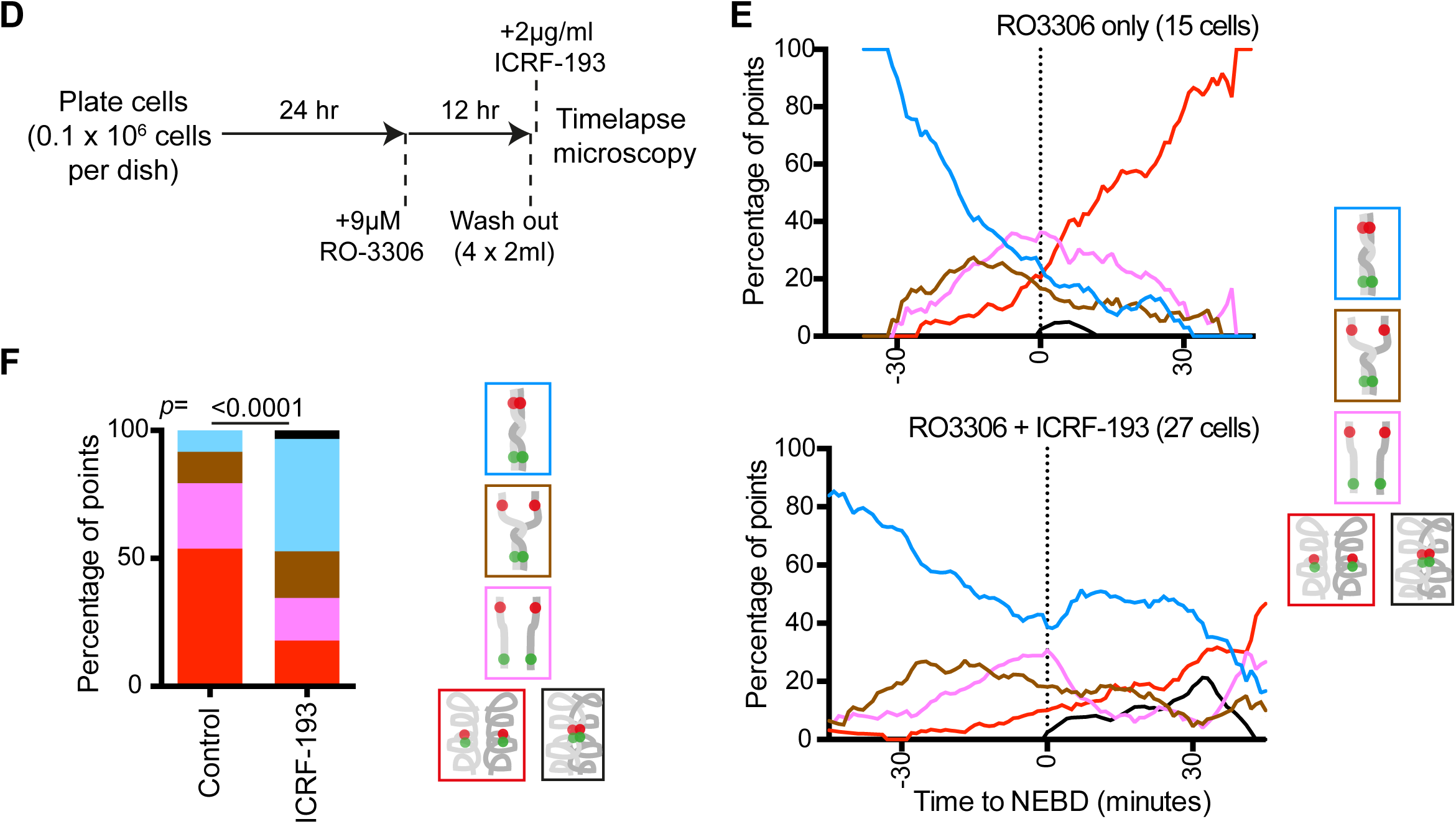
related to Figure 4. (A) Experimental procedure outline. Cells were synchronised in early S-phase using a double thymidine block. Released cells were treated with MK1775 (WEE1 inhibitor) 1 hour before, and ICRF-193 (topo II inhibitor) 5–7 mins before, time-lapse imaging. (B) Behaviour of the fluorescent reporter as observed in individual live-cells, plotted across the Y-axis. TT75 cells were arrested by a double thymidine block and subsequently released from the block. 8–10 hr after the release cells were treated with MK1775 and ICRF-193 as described in part (A), images were taken every minute and, at each time point, the configuration of the reporter was determined and represented using the colour-codes depicted in Figure 1(C) and (D). Data from individual cells was aligned according to Figure 1(E). White spaces represent time points where configuration could not be determined. (C) Evaluating specificity of the black state. With WAPL depletion and with ICRF-193 treatment, we often observed persistent (at least for 5 consecutive time points) co-localisation of all four fluorescent dots, following NEBD (Figure 2E, right bottom; Figure 3B, right; Figure S4E, bottom). We interpreted it as a “non-resolved and compacted” state and defined it as a black state. The black state was specific to WAPL depletion and ICRF-193 treatment, as it was very rarely observed in other conditions after NEBD. Next we evaluated how often the configuration corresponding to the black state appeared before NEBD. For this, we applied the criteria for the black state (see Experimental procedures) to the time points before NEBD, in the data sets for control siRNA, WAPL siRNA (Figure 2E), control treatment and ICRF-193 treatment (Figure 4B). The configuration corresponding to the black state was observed in these conditions before NEBD, but only at a low percentage, as shown here. We interpret it as fortuitous continuous co-localisation of red and green fluorescent dots (on non-resolved sister chromatids), since chromosome compaction is not substantial before NEBD (see the red “compacted” state in various conditions). Based on this interpretation, the configuration corresponding to the black state before NEBD was included in the blue “non-resolved” state in all analyses in figures, except for the figures shown here. Collectively these data show that frequent appearance of black states (i.e. persistent co-localisation of all four fluorescent dots) is specific to a) WAPL depletion and ICRF-193 treatment and b) post-NEBD. (D) Experimental procedure outline. Cells were synchronised at the G2-M boundary using RO-3306 (CDK1 inhibitor). Released cells were treated with ICRF-193 5-7 mins before time-lapse imaging. (E) Behaviour of the fluorescent reporter observed in control cells or cells treated with ICRF-193 live-cells. TT75 cells were synchronised as in part (C) and immediately after the release images were taken every minute and, at each time point, the configuration of the reporter was determined and represented using the colour-codes depicted on right. Data from individual cells was aligned according to NEBD and proportions of each configuration at each time point was determined with smoothing applied by calculating the rolling mean proportion across 9 timepoints. (F) The change in fluorescent reporter configuration following NEBD was scored according to the pipeline shown in Figure 4(E), left. The proportion of each state during the 20 minutes following NEBD for the selected cells is shown for control or ICRF-193 treated cells. *p* value was obtained by *chi*-squared test and the number of analysed time points were 79 and 116 for control and ICRF-193 treated cells, respectively.

**Figure S5:**
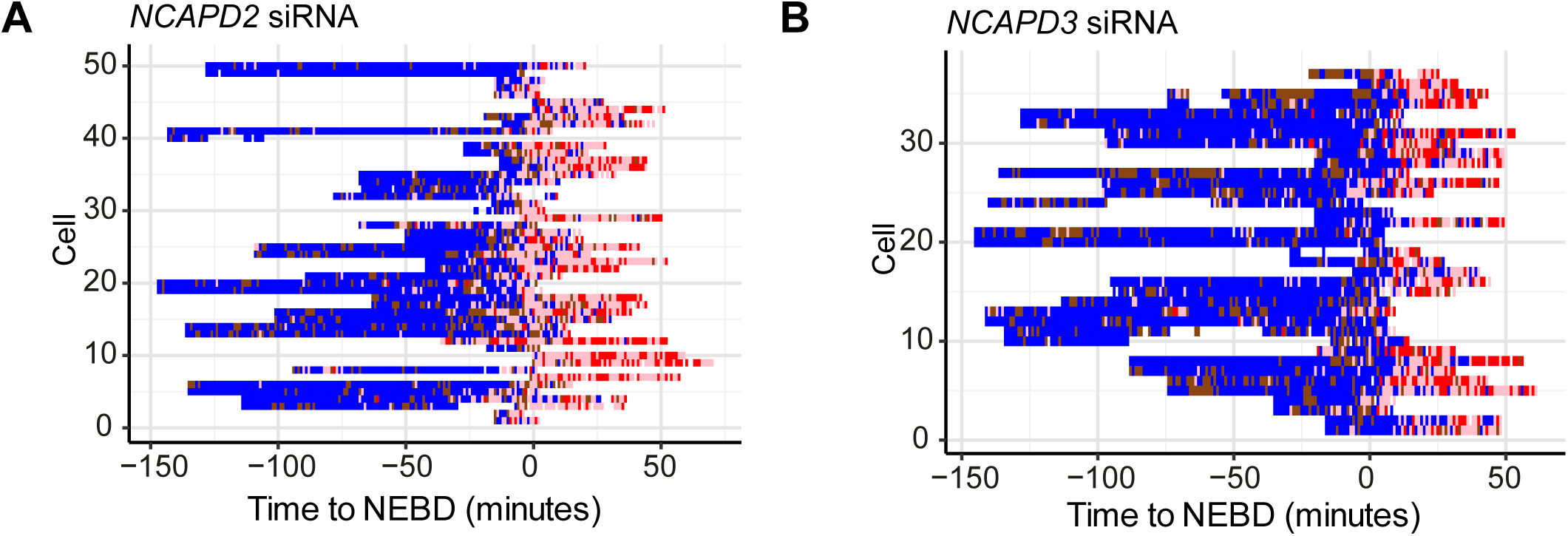
(related to Figure 5) (A) and (B) Behaviour of the fluorescent reporter as observed in individual live-cells, treated with NCAPD2 (condensin I) or NCAPD3 (condensin II) siRNA, plotted across the Y-axis. TT75 cells were treated with siRNA, arrested by a double thymidine block and subsequently released from the block. 8–10 hr after the release, images were taken every minute and, at each time point, the configuration of the reporter was determined and represented using the colour-codes depicted in Figure 1(C) and (D). Data from individual cells was aligned according to Figure 1(E). White spaces represent time points where configuration could not be determined.

**Figure S6:**
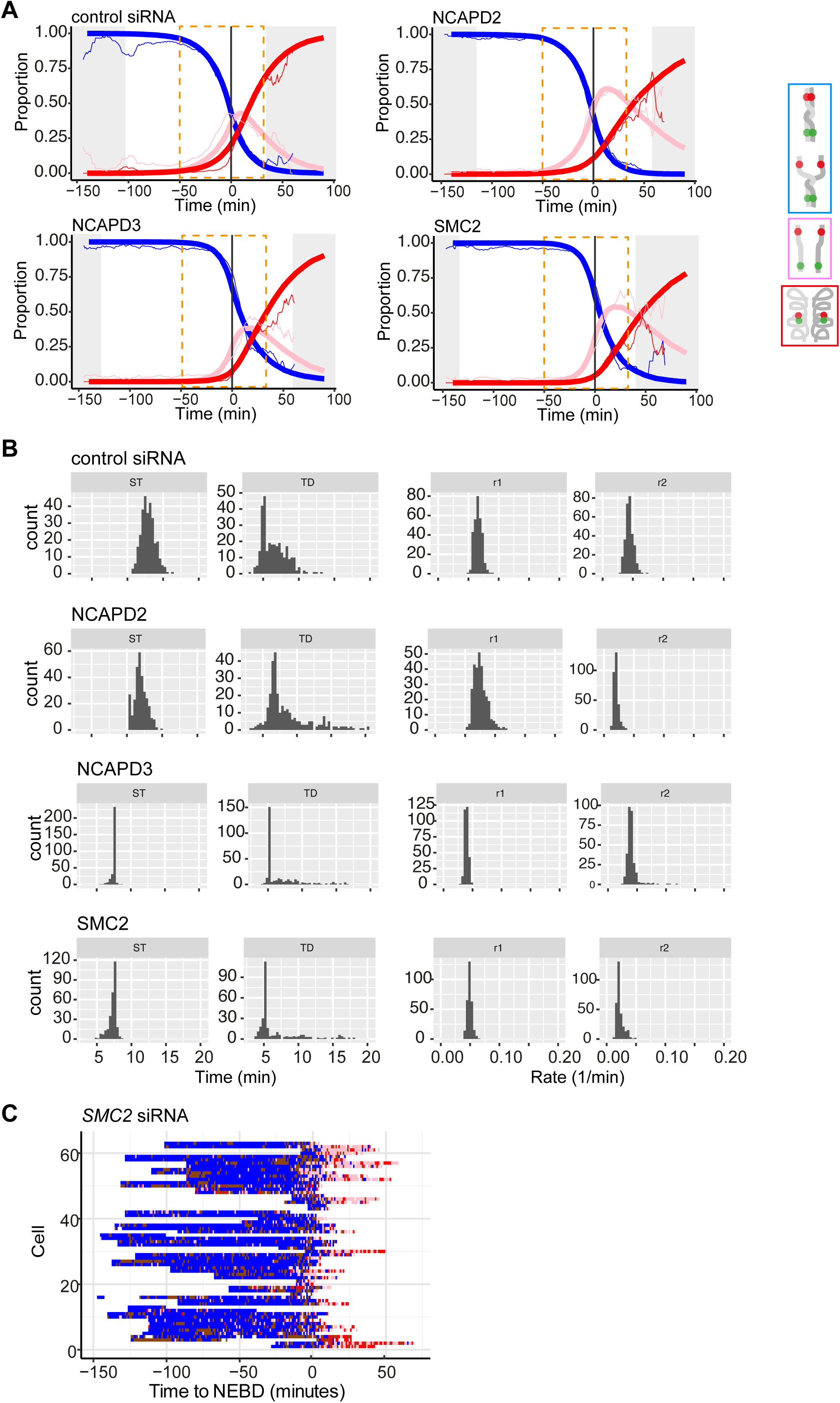
(related to Figure 6) (A) Control, NCAPD2, NCAPD3 and SMC2 siRNA treated cells data (thin curves) with their best-fitting models (thick curves). The curves represent the proportion of each configuration in the fluorescent reporter system at each time point. The data curves were smoothed by a running mean with window size of 5 minutes for presentation only (that is, fitting was performed on unaltered data). The colour codes of lines are in the legend. The grey shaded area is as in Figure 1(F). The orange dotted box indicates the region over which the mean-square difference between the data and the model was calculated. (B) Results of a bootstrap performed to assess model parameter uncertainties for control, NCAPD2, NCAPD3 and SMC2 siRNA treated live-cells (TT75). In each bootstrap data were sampled with replacement and the best fit was found. 300 bootstraps were performed to create the distributions shown. See Experimental procedures for more details. (C) Behaviour of the fluorescent reporter as observed in individual live-cells, treated with SMC2 (condensin I and II) siRNA, plotted across the Y-axis. TT75 cells were treated with siRNA, arrested by a double thymidine block and subsequently released from the block. 8–10 hr after the release, images were taken every minute and, at each time point, the configuration of the reporter was determined and represented using the colour-codes depicted in Figure 1(C) and (D). Data from individual cells was aligned according to Figure 1(E). White spaces represent time points where configuration could not be determined.

